# Actinomycin D Drives RNA-Binding Proteins into Dynamic Cytoplasmic Granules

**DOI:** 10.64898/2026.08.18.745449

**Authors:** Aydan Torun, Hoşnaz Tuğral Dunuroğlu, Ekinsu Gürsöz, Leman Nur Nehri, Nurhan Özlü, Erol Yıldırım, Sreeparna Banerjee

## Abstract

Actinomycin D (Act D) is a global transcriptional inhibitor widely used in research and clinical practice; however, its effects on RNA-binding protein (RBP) dynamics remain poorly understood. Analysis of an RNA-seq dataset from Act D-treated HeLa cells revealed a compensatory stress response enriched in RNA metabolism, processing, and translation. Here, we investigated the effects of Act D on the subcellular localization of RBPs using HuR as a model mRNA stabilizing RBP. Short-term Act D treatment markedly increased cytoplasmic HuR localization in HCT116 and HeLa cells where the protein is known to be active. Analysis of known pathways regulating HuR nucleocytoplasmic translocation did not fully explain this redistribution, suggesting alternative mechanisms.

To identify proteins proximal to HuR following Act D treatment, we performed TurboID labeling followed by LC-MS/MS in HCT116 cells. Several proteins involved in RNA regulation were identified. Probabilistic modeling highlighted FUS, an RBP with established roles in phase-separated granule dynamics, as a candidate proximal protein. The Act D-dependent interaction between HuR and FUS was interrogated using molecular dynamics simulations and validated with proximity ligation assays. Furthermore, increased cytoplasmic localization of RBPs following Act D treatment was accompanied by formation of granular structures that were relatively fluid and could be disrupted by hypotonic shock.

Collectively, our findings demonstrate that Act D induces cytoplasmic redistribution of multiple RBPs and their sequestration into dynamic granular structures, revealing a previously unrecognized cellular response to transcriptional inhibition.

**Graphical Abstract:** Act D induced cytoplasmic re-localization of HuR along with FUS and other RBPs in dynamic, hypotonic shock-sensitive granular structures.

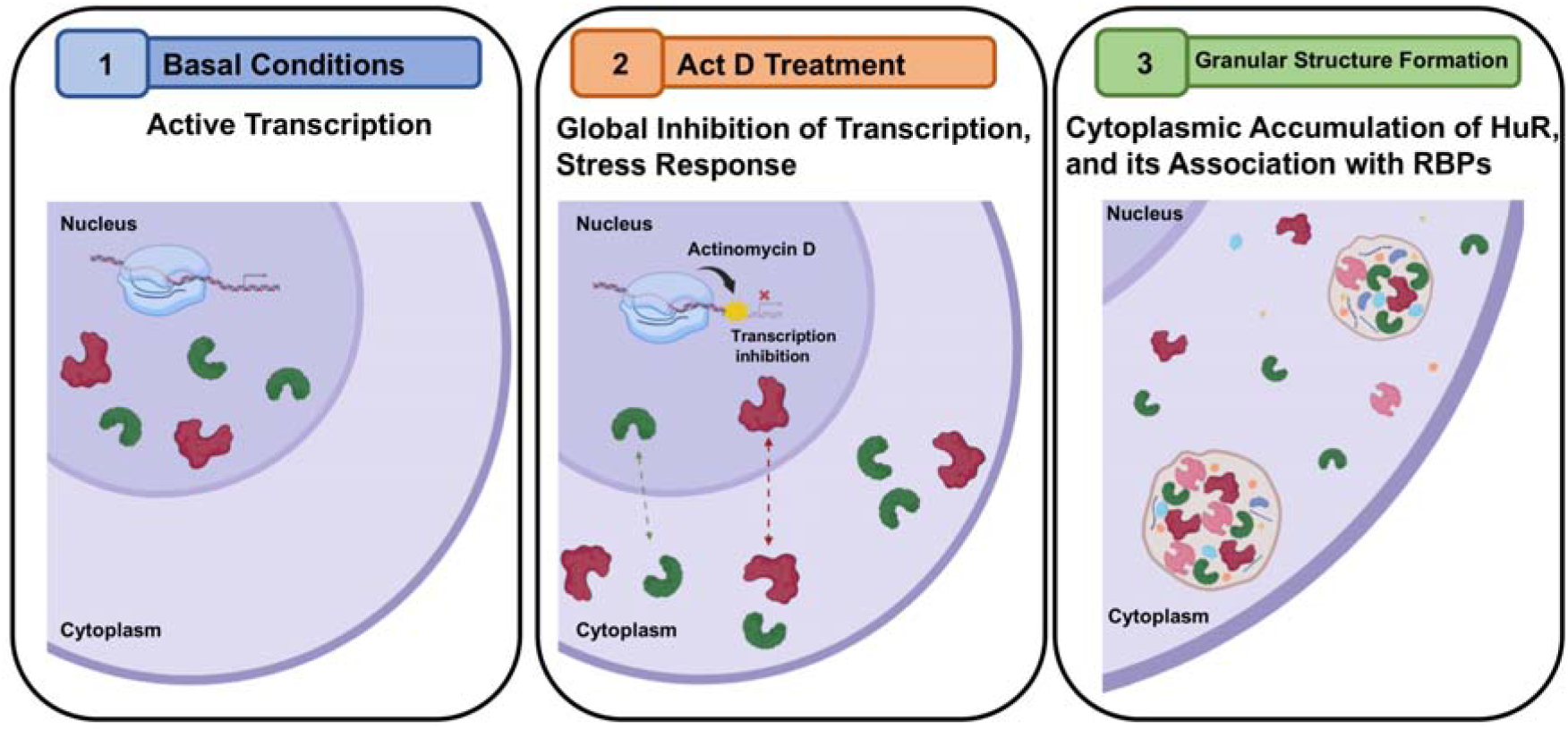

## 1 Introduction

Cells are continuously exposed to a variety of environmental and intracellular stressors that can induce a coordinated gene regulatory program. A substantial and rapid component of stress adaptation occurs at the post-transcriptional level via RNA-binding proteins (RBPs) (1, 2). Upon exposure to oxidative stress, heat and cold shock, hypoxia, ultraviolet light (UV) etc., signaling cascades activate RBPs to modulate RNA stability, translation efficiency, and localization to cytoplasmic condensates (3). Therefore, RBPs are critical determinants of RNA fate and function through dynamic changes in their activity, localization, and interactions (1).

RBPs also regulate multiple aspects of RNA metabolism, including alternative splicing, polyadenylation, stability, localization, translation, and miRNA processing, and alterations in these functions can contribute to different diseases including cancer (4). Consistent with this, accumulating evidence indicates that RBPs play critical and versatile roles across multiple Hallmarks of Cancer (5), positioning them as emerging targets in cancer research and therapy (6, 7).

HuR (*ELAVL1*) is a ubiquitously expressed RBP that plays a central role in post-transcriptional gene regulation, particularly during cellular stress. Unlike other members of the ELAVL family which are largely restricted to neuronal tissues, HuR is broadly expressed and increases the stability and translation of target mRNAs by binding AU-rich elements (ARE) within their 3′ untranslated regions (8). HuR contains three conserved RNA recognition motifs (RRM) linked by a highly basic, intrinsically disordered hinge region that harbors the HuR nucleocytoplasmic shuttling sequence (HNS), enabling its dynamic transport between the nucleus and cytoplasm (9, 10). Cytoplasmic accumulation of HuR is a critical determinant of its function and is regulated by multiple signaling pathways, including AMPK (11), p38/MK2 (12), various PKC isoforms (13, 14), and Chk2 (15), which modulate the localization of the protein through post-translational modifications. Consistent with its role in stress responses and mRNA regulation, HuR has been implicated in a range of diseases, including cardiometabolic (16), immune (17), neurological disorders (18), and cancer (19). Moreover, elevated cytoplasmic localization of HuR has been correlated with tumor grade and poor survival outcomes across various cancer types (17). Recent studies further highlight the importance of HuR shuttling in cell fate decisions, demonstrating that impaired cytoplasmic translocation can affect its cleavage during apoptosis and promote cell survival, underscoring its potential relevance in cancer biology (20, 21).

The formation of phase separated granules or biomolecular condensates from proteins and nucleic acids has been widely accepted in recent years as a stress response to changing environmental and physiological conditions (22, 23). RBPs are critical components of condensates as they harbor intrinsically disordered region (IDR) that enable them to form low-specificity, multivalent interactions with other proteins and nucleic acids, thereby triggering their compartmentalization into condensates (24, 25). HuR is frequently identified as a component of condensates including stress granules (26, 27), and gel-like TIS11B granules (28), and was reported to be compartmentalized on microtubules along with RBPs like FUS, TDP-43, and G3BP1 (29).

Transcriptional inhibition with the Topoisomerase I inhibitor Camptothecin was reported to induce the accumulation of the genome integrity protein SLX4 in nuclear condensates (30). More recently, condensates induced by transcription inhibition (CITIs) were identified in cells undergoing transcriptional inhibition due to UV, heat shock or chemotherapeutic drugs (31). Actinomycin D (Act D) is an antibiotic that inhibits all three eukaryotic RNA polymerases (32) and is used therapeutically for Wilm’s tumor, Ewing’s sarcoma and rhabdomyosarcoma (33).

The current study was based on an analysis of a publicly available RNA-seq dataset (GSE198178) (34) which revealed that treatment of HeLa cells with Act D led to the enrichment of GO terms related to RNA biology. Since Act D is commonly used in laboratories for studying post-transcriptional regulation and half-life of mRNA transcripts, we hypothesized a role of Act D in the activity on RBPs. Focusing on HuR, we observed a remarkable and rapid cytosolic translocation of HuR upon treatment with Act D. This translocation was not dependent on known pathways such as Chk2 activity or DNA damage. Additionally, HuR protein formed typical aggregates with other RBPs that were dynamic and the constituent proteins were retained in the cytoplasm up to 16h after the washout of Act D. These aggregates could be disassembled under hypotonic conditions. Overall, our data shows for the first time that phase separation and aggregation of RBPs in response to treatment of cells with Act D and should be considered when evaluating mRNA half-life.

## 2 Materials and Methods

### 2.1. RNA-seq Analysis

Raw FASTQ files from the publicly available RNA-seq dataset GSE198178 (34) were uploaded to the Galaxy platform (use.galaxy.eu) for differential gene expression analysis. The dataset consisted of single-end sequencing reads. Adapter sequences and low-quality reads were removed using Trimmomatic (35). The quality control of the trimmed reads was carried out using FastQC. The cleaned reads were aligned to the human reference genome (hg19) with HISAT2 (36), which provided the resulting alignments as BAM file outputs. Gene expression was determined with the read counting program featureCounts (37). Gene names were obtained with a generic set of identifiers via the annotateMyIDs tool. Differential expression analysis was carried out using the voom-limma method in the *limma* package. Library sizes were normalized using the trimmed mean of M-values (TMM) normalization method. Three replicates were included for each experimental group. Differentially expressed genes (DEGs) were identified by comparing 1 h, 2 h, and 4 h Act D treated samples with the untreated control group (0h). Statistical significance was determined using adjusted p-values (pAdj), and genes with p(Adj) < 0.05 were considered significantly differentially expressed. Gene expression changes were represented as log_2_ fold change (log_2_FC), and genes with log_2_FC > 0 were classified as upregulated. Significantly upregulated genes from each treatment time point were compared to identify common genes among all three groups. These commonly upregulated genes were subsequently analyzed using the STRING database for Gene Ontology enrichment analysis. Enrichment in STRING is reported as Strength (log_10_(observed/expected)); therefore, fold enrichment (observed/expected number of genes associated with a given GO term) was calculated as 10^Strength for visualization.

### 2.2. Cell Culture and treatments

HCT116 cell line was purchased from DKFZ (Heidelberg, Germany) and the cells were grown in RPMI 1640 without phenol red (VivaCell Biosciences, Shanghai, China) supplemented with 10% FBS, 2mM L-glutamine and 1% penicillin/streptomycin. The HeLa cell line (ATCC) was grown in Eagle’s minimum essential medium (EMEM) (VivaCell Biosciences) supplemented with 10% fetal bovine serum (FBS) (Capricorn Scientific, Germany), 2 mM L-glutamine (Sartorius, Germany), 1 mM sodium pyruvate (Sartorius), 0.1 mM non-essential amino acids (VivaCell Biosciences) and 1% penicillin/streptomycin (Sartorius). The cells were incubated at 37 °C with 5% CO_2_ and treated with 2.5 μg/mL Plasmocin® (Invivogen, France) for the prevention of mycoplasma contamination. The cells were routinely tested for mycoplasma contamination (38). The reagents used in this study include Act D (Tocris, 1229), -Amanitin (Cayman, 17898), Camptothecin (Cell Signalling Technology, 13637), BML277 (Selleckchem, S8632), Leptomycin B (Sigma-Aldrich, L2913), and Biotin (Sigma-Aldrich B4639), RNase A (Thermo Fisher Scientific, EN0531).

### 2.3. Plasmids, Constructs & Transfections

Full length HuR gene (HuR-FL) and the HuR gene lacking the HNS (ΔHNS) were cloned into a pEGFP-C1 vector using *BglII* and *SalI* restriction sites at the 5’ and 3’ends, respectively. Myc-tagged HuR was cloned into pcDNA3.1-TurboID using the *XbaI* and *NotI* restriction sites. The cloned DNA fragments were amplified by using specific primers with Q5 DNA Polymerase (NEB). A pGEX-6P-1 ELAVL1 (Dundee University MRC PPU Reagents & Services) expression vector was used as the template. The vectors and the inserts were cut with the indicated restriction enzymes and ligated using T4 DNA Ligase (NEB) O/N at 16 °C. The ligation products were transformed into *E.coli* NEB Stable competent cells and the colonies were screened for the presence of the insert via PCR. Plasmids isolated from the positive colonies were analyzed via next generation sequencing for the confirmation of successful cloning. Transient transfections of the plasmids were carried out for 24 hours using Xtreme Gene Transfection Reagent (Sigma-Aldrich) according to the manufacturer’s instructions.

### 2.4. Protein Isolation and Western Blot Analysis

Total proteins were isolated using M-PER Mammalian Protein Extraction Reagent (Thermo Fisher Scientific) supplemented with protease and phosphatase inhibitors (Roche, Germany). For cytoplasmic and nuclear fractionation, cell pellets were lysed in hypotonic buffer (10 mM HEPES (pH 7.5), 4 mM sodium fluoride, 10 μM sodium molybdate, 0.1 mM EDTA, and protease/phosphatase inhibitors) followed by addition of 1.5% NP-40 (AppliChem). The supernatant was collected as the cytoplasmic fraction after centrifugation. The nuclear pellet was lysed by sonication in RIPA buffer containing protease and phosphatase inhibitors. After centrifugation, the supernatant was collected as the nuclear fraction.

Protein concentrations were measured using Coomassie Protein Assay Reagent (Thermo Fisher Scientific) according to the manufacturer’s instructions. Equal amounts of proteins within each fraction were separated on 10% SDS-PAGE gels and transferred onto PVDF membranes (Roche). Membranes were blocked with 5% skim milk in TBS-T and incubated with primary antibodies overnight at 4 °C, followed by incubation with HRP-conjugated secondary antibodies for 1 h at room temperature. Protein bands were visualized using Clarity ECL substrate (Bio-Rad) on a ChemiDoc MP Imaging System (Bio-Rad). Band intensities were quantified using Image Lab software and normalized to housekeeping proteins. Antibodies used in this study are listed in Supplementary Table 1.

### 2.5. TurboID Assay

HCT116 cells were transfected with either empty vector (pcDNA3.1-TurboID-HA) or HuR-TurboID constructs. Twenty-four hours post-transfection, the cells were treated with 10 μg/mL Act D and incubated with 50 μM biotin and 1 mM ATP for 10 or 30 min. Following the optimization of lysis conditions, cells were lysed in buffer containing 50 mM Tris-HCl (pH 7.4), 500 mM NaCl, 0.4% SDS, 5 mM EDTA, 2% Triton X-100, 1 mM DTT, and protease inhibitors, followed by sonication and centrifugation. Protein concentrations were determined using Pierce BCA Protein Assay Kit (Thermo Fisher Scientific).

For streptavidin pull-down, 6 mg of protein lysate was incubated overnight at 4°C with streptavidin magnetic beads (NEB). Bead-bound proteins were washed with SDS-, deoxycholate-, NP-40-, and Tris-based wash buffers. A portion of the samples was analyzed by SDS-PAGE and Western blotting to confirm biotinylation efficiency, while the remaining samples were subjected to LC-MS/MS analysis at the Koç University Proteomics Facility (KUPAM) sing an UltiMate 3000 RSLCnano reversed-phase chromatographic platform (Thermo Fisher Scientific) coupled to a Q-Exactive quadrupole-Orbitrap mass spectrometer (Thermo Fisher Scientific). Raw MS data were processed using Proteome Discoverer (version 2.3) with the Sequest HT search engine. The search was subjected against the UniProt Homo sapiens database (21,039 protein entries) (39).

TurboID experiments were performed in two independent biological replicates, each analyzed with three LC-MS/MS technical replicates. Proteins common across technical and biological replicates were identified, and proteins specifically enriched in Act D-treated HuR-TurboID samples were determined using the Bioinformatics & Evolutionary Genomics Venn diagram tool. Protein interaction network and gene ontology enrichment analyses were performed using STRING. For visualization of GO enrichment, fold enrichment was calculated from the STRING Strength metric (10^Strength).

### 2.6. Immunofluorescence and Proximity Ligation Assay

Following treatments or transfections, cells grown on coverslips were fixed with 4% paraformaldehyde for 15 min at room temperature, permeabilized with 0.1% Triton X-100, and blocked with 5% BSA. Cells were incubated with primary antibodies diluted in 1% BSA, followed by incubation with fluorescence-conjugated secondary antibodies. Nuclei were counterstained using ProLong Diamond Antifade Mountant with DAPI (Thermo Fisher Scientific). Fluorescence images were acquired using either a Zeiss Axio Imager M2 or Olympus BX43 microscope.

For proximity ligation assays (PLA), HeLa cells were seeded on glass coverslips and treated with Act D for 3 h prior to fixation with 4% paraformaldehyde. PLA was performed using the Duolink In Situ PLA kit according to the manufacturer’s instructions. Images were acquired using an Olympus BX43 fluorescence microscope at 100× magnification.

### 2.7. Molecular Docking and Dynamic Simulations

Protein-protein docking was carried out to investigate the interaction interface between HuR and FUS. After docking modelling, two molecules of Act D were manually positioned within the interaction interface region to generate the HuR-FUS-2ActD complex. Molecular dynamics simulations were subsequently carried out for 3 ns using the COMPASS II force field under the NPT ensemble conditions. Mean square displacement (MSD) analysis was performed to evaluate the dynamic behavior and structural stability of the complexes during the simulation period. Binding energy calculations were carried out by subtracting the energies of their individual components from the total energy of the complexes to understand the interaction strength.

### 2.8 Probabilistic modeling

A probabilistic model was developed to integrate experimentally derived protein features associated with HuR into a unified computational framework and provide a systems-level interpretation of HuR-associated granule assembly. Proteins identified by the TurboID assay were categorized based on protein-protein interaction (PPI) scores, condensate localization, intrinsic disorder region (IDR) annotations, and isoelectric point (pI) values (Figure S13). To construct the model parameters, PPI scores were treated as continuous variables, whereas condensate localization and IDR annotations were encoded as binary variables (0 or 1).

Proteins lacking information for any of the four features were excluded from downstream analyses, resulting in a final dataset of nine proteins.

The normalized isoelectric point (p*I)* was calculated using min-max normalization:

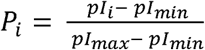

An Assembly Score was calculated as a weighted linear combination of four biological features:

A_i_=*aI_i_*+Βc_i_*+γD_i_+δP_i_; where A_i_=the Assembly Score of protein i;I_i =_*PPI score ;*condensate* localization (0 or 1); =intrinsic disorder region (IDR) annotation (0 or 1) and P_i_=the normalized isoelectric point.

Because the relative contributions of the four biological features were unknown, the weighting coefficients (α,β,γ,δ) were randomly sampled from a Dirichlet(1,1,1,1) distribution such that their sum equaled 1. A total of 10,000 independent Monte Carlo simulations were performed. For each iteration, a new Assembly Score was calculated and subsequently interpreted as the transition probability governing granule assembly (), representing the probability of progressing between consecutive cellular states.

A transition probability matrix was defined to describe movement between consecutive cellular states. At each simulation step, proteins either remained in their current state with probability (1-p) or transitioned to the next assembly state with probability . Direct transitions between non-adjacent states were not permitted, and the Stable Granule state was treated as an absorbing state. Each simulation was initialized using s_0_ =[1 0 0 0], indicating that all proteins initially resided in the nucleus. For each protein, the mean Assembly Score and the mean Stable Granule probability were calculated across all Monte Carlo simulations:

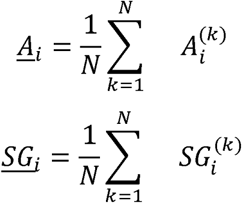

where N=10,000; *A*^(*k*)^_*i*_ = Assembly Score of protein *i* in Monte Carlo iteration *k*, and *SG*^(*k*^)_*i*_ the corresponding Stable Granule probability.

Finally, sensitivity analysis was performed by correlating the randomly sampled feature weights with the predicted Stable Granule probabilities to estimate the relative contributions of each biological feature to granule assembly. The analysis pipeline is shown in Figure S14.

### 2.9. Statistical Analysis

Statistical analyses were performed using GraphPad Prism. Statistical significance was assessed using either Student’s t-test or one-way ANOVA, as appropriate. A p value < 0.05 was considered statistically significant. All analyses were conducted using data obtained from at least 2-3 independent biological replicates.

## 3 Results

### 3.1 Global Transcription Inhibition Increased the Cytosolic Translocation of HuR

Act D is known to globally inhibit transcription. Examination of RNA-seq data (GSE198178) from Act D treated (5μM for 1h, 2h, 4h) HeLa cells, however, revealed that while most of the mRNAs were downregulated as expected, upregulation of some mRNAs could be observed (Figure 1A). Gene ontology (GO) analysis of the significantly and consistently upregulated transcripts at all time points revealed the enrichment of genes involved in ribosome biogenesis, translation, rRNA processing, and RNA-binding functions as well as non-membrane bounded organelle assembly and post-transcriptional regulation (Figure 1B, Figure S1A−D). This suggests that treatment of HeLa cells with Act D could induce a compensatory stress response for RNA metabolism and potentially altering the stability of the transcripts.

**Figure 1.**
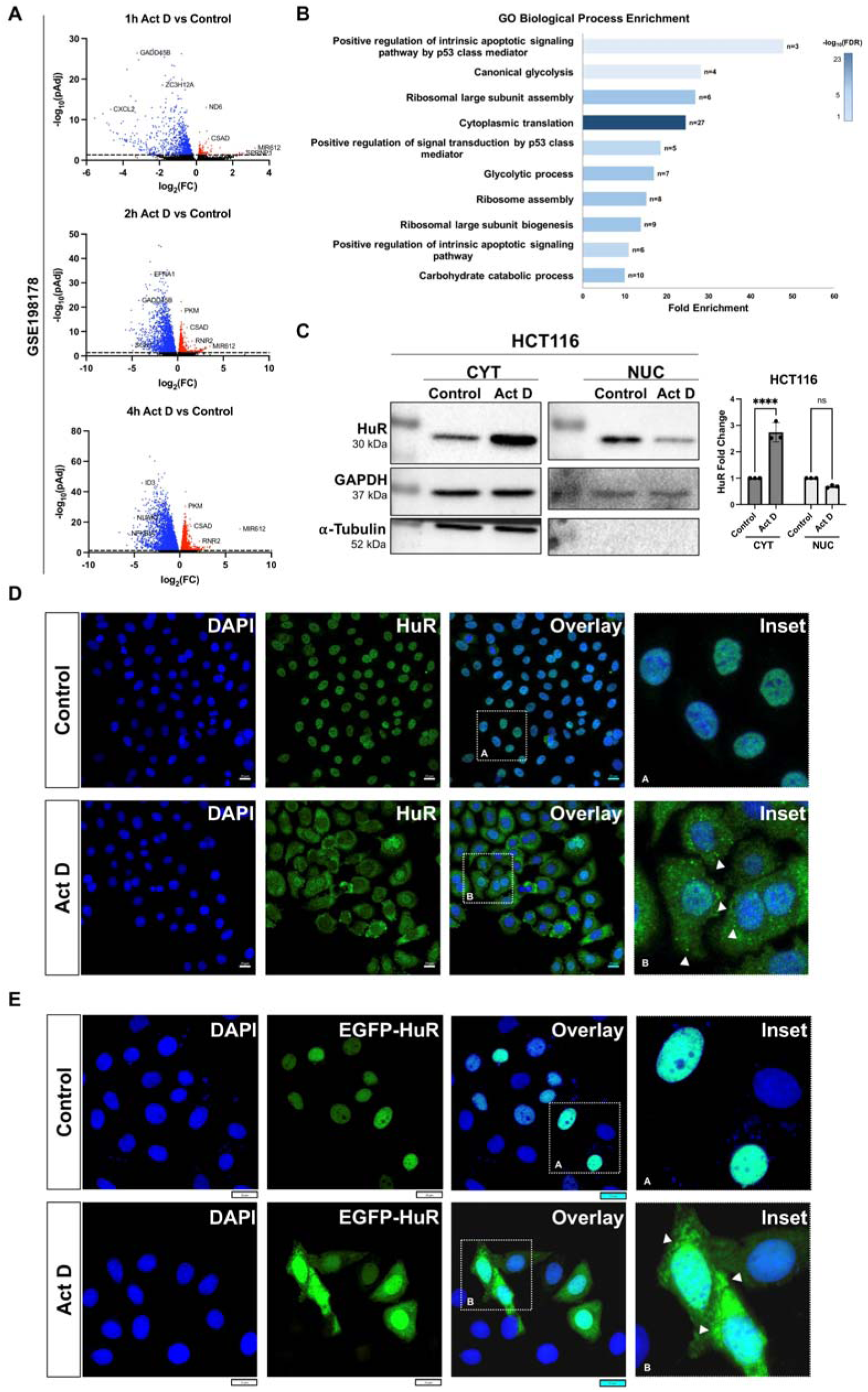
Actinomycin D-Induced Cytoplasmic Translocation of HuR in HCT116 and HeLa Cells. (A) The RNA-seq dataset GSE198178, in which Hela cells were treated 5 μM Actinomycin D (Act D) for 1h, 2h or 4h, was analyzed via Galaxy platform. Differentially expressed genes at each time point were graphed. Significantly upregulated genes are shown in red while significantly downregulated genes are shown in blue. (B) Gene ontology enrichment analysis for biological processes was performed on significantly upregulated genes (p<0.05 and log_2_(FC)>0) using STRING. Top 10 enriched biological process terms are ranked by fold enrichment (10^Strength, where Strength=log_10_(observed/expected), as reported by STRING). Bar length represents fold enrichment, bar color shows –log_10_(FDR), and labels (n) indicate the observed number of genes associated with each GO term. (C) HCT116 cells were treated with 10 μg/mL Act D for 3 h. Cytoplasmic (CYT) and nuclear (NUC) fractions were isolated, and HuR protein levels in each fraction were analyzed by immunoblotting. Representative blots are shown. Densitometry quantification of HuR levels in Act D-treated cells relative to control cells (serum free medium, SFM) was performed separately for cytoplasmic and nuclear fractions. Data represent three independent biological replicates. Statistical significance was assessed using an unpaired t-test (****p < 0.0001; ns, not significant). GAPDH and α-tubulin were used as loading controls. (D) HeLa cells were treated with 10 μg/mL Act D for 3 h and processed for immunofluorescence staining using antibodies against HuR, with nuclei counterstained by DAPI. Representative images were acquired using a Zeiss Axio Imager M2 microscope. Scale bar, 20 μm. (E) HeLa cells transfected with the pEGFP-c1-HuR Full-Length construct (EGFP-HuR) were treated with 10 μg/mL Act D for 3 h. Following fixation, nuclei were stained with DAPI, and GFP fluorescence was used to visualize overexpressed HuR. Images were acquired using an Olympus BX43 microscope. Scale bar, 20 μm. Insets show magnified views of the indicated regions.

As a model RBP, we selected HuR which is known to translocate to the cytoplasm upon stimulation, bind to the 3’UTR of target mRNAs and stabilize them. We observed that the treatment of HCT116 (Figure 1C) or HeLa (Figure 1D) cells with 10 μg/mL (approximately 8 μM) Act D for 3 h resulted in an increase in the cytoplasmic levels of the endogenous HuR protein. Immunofluorescence staining showed a remarkable translocation of HuR to the cytoplasm when HeLa cells were treated with Act D (10 μg/mL, 3h) (Figure 1D). Moreover, we observed an increase in the total protein levels of HuR with Act D treatment (10 μg/mL, 3h) in both cell lines (Figure S2).

To evaluate whether ectopically expressed HuR protein could also be translocated to the cytoplasm with Act D treatment, we transfected HeLa cells with HuR tagged to EGFP. While the exogenously expressed EGFP-HuR protein was localized in the nucleus as expected (Figure 1E), Act D treatment (10 μg/mL, 3h) led to the cytoplasmic localization of the overexpressed protein as well (Figure 1E). Treatment of the cells with α-Amanitin (α-AMA), a selective inhibitor of RNA Polymerase II and III, also led to an increase in the cytoplasmic levels of HuR, but only when the cells were treated for over 16 h (Figure S3A, B). These data suggest that Act D-induced stress in both cell lines triggered a rapid movement of HuR (both endogenous and overexpressed) to the cytoplasm.

### 3.2 Evaluation of Known Regulatory Pathways in Actinomycin D-Induced HuR Cytoplasmic Localization

We next sought to explore the mechanisms for the cytoplasmic localization of HuR upon Act D treatment. Various pathways and mechanisms are known to regulate the cytoplasmic translocation of HuR including the p38-MK2 pathway (12) and the DNA damage response kinase Chk2 (15). We observed that Act D did not lead to the phosphorylation of either p38 or MK2 proteins (data not shown); these pathways were not evaluated further. An increase in the phosphorylation of Chk2 was observed with Act D treatment (10 μg/mL, 3 h) (Figure 2A, B, Figure S4A, B). Co-treatment of the cells with the Chk2 inhibitor BML277 and Act D led to a decrease in the phosphorylation of Chk2, as expected, but did not inhibit the cytosolic translocation of HuR. This suggests Act D treatment led to the activation of DNA damage response and Chk2 phosphorylation; however, this pathway was not responsible for the cytoplasmic accumulation of HuR. We next investigated whether an alternative DNA damage inducing agent could also increase the cytoplasmic translocation of HuR. For this, we treated the cells with the DNA topoisomerase I inhibitor Camptothecin (CPT). Although Chk2 was activated in the presence of CPT, we did not observe any cytoplasmic translocation of HuR (Figure S5A, B). This suggested that mechanisms other than DNA damage response could be responsible for the cytoplasmic accumulation of HuR upon Act D treatment.

**Figure 2.**
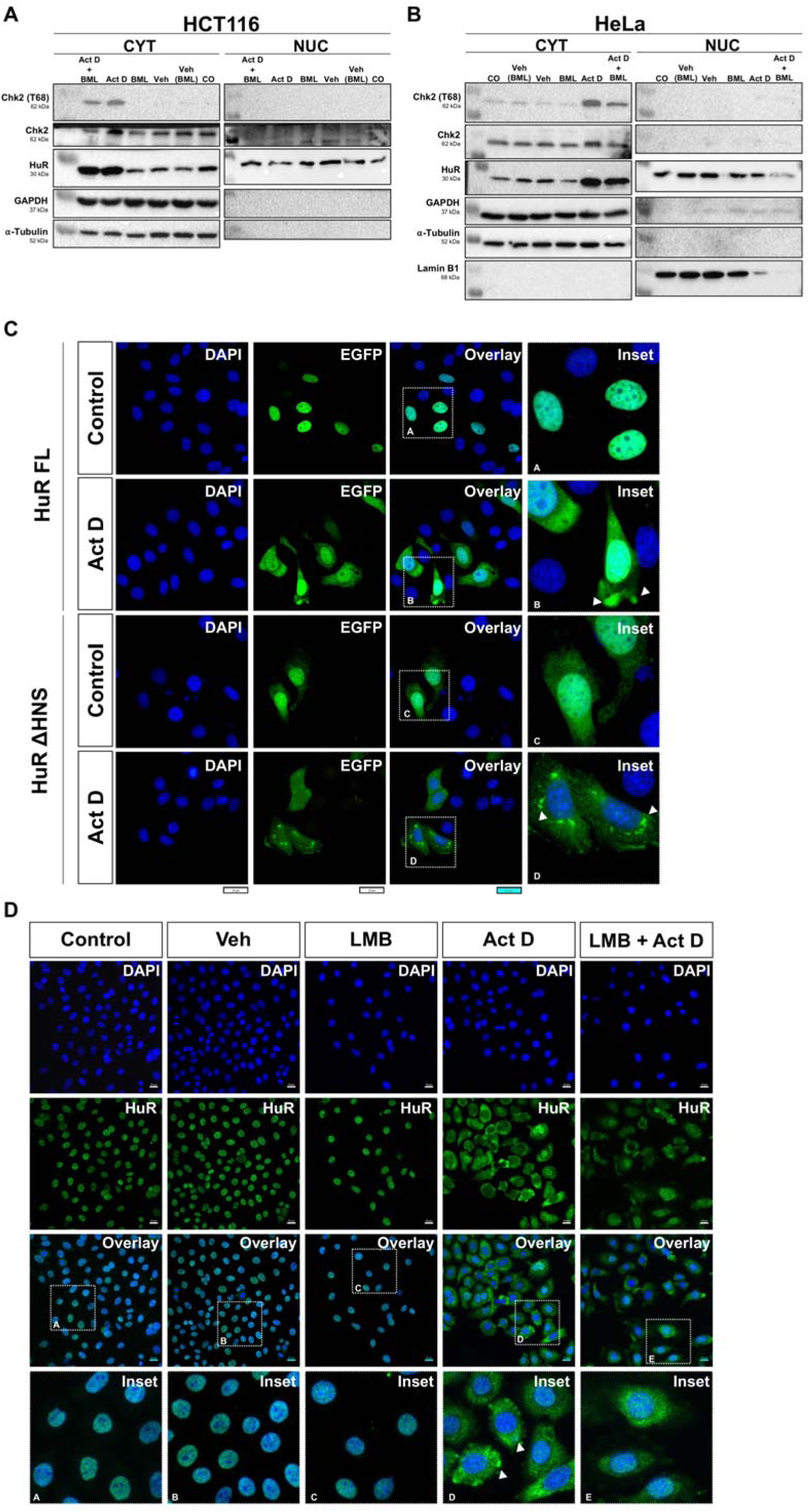
Robust Cytoplasmic HuR Accumulation Persists Despite Inhibition of Known Regulatory Mechanisms(A-B) HCT116. (A) and HeLa (B) cells were treated with 10 μM Chk2 inhibitor BML277 (BML) for 48 h, in the presence or absence of Actinomycin D (Act D). Cells were subsequently fractionated into cytoplasmic (CYT) and nuclear (NUC) compartments. GAPDH and α-tubulin were used as loading controls for cytoplasmic fractions, while Lamin B1 served as a nuclear loading control. For statistical analysis, two independent biological replicates were performed for HCT116 cells and three independent biological replicates for HeLa cells. Densitometric quantifications of cytoplasmic levels of HuR and Chk2 (T68) are presented below the corresponding blots. Statistical significance was determined using one-way ANOVA followed by Tukey’s post hoc test (*p < 0.05, **p < 0.005, ****p < 0.0001; ns, not significant). (C) HeLa cells were transfected with pEGFP-c1-HuR Full-Length (HuR FL) or pEGFP-c1-ΔHNS (HuR ΔHNS) constructs for 24 h. Following transfection, cells were either treated with 10 μg/mL Actinomycin D (Act D) for 3 h or left untreated as controls. GFP fluorescence, representing the expressed HuR variants, was imaged using an Olympus BX43 microscope. (D) HeLa cells were treated with Leptomycin B (LMB) in the presence or absence of Actinomycin D (Act D). Following treatment, cells were fixed and immunofluorescence staining was carried out using an anti-HuR antibody and an Alexa Fluor 488-conjugated secondary antibody. Nuclei were counterstained with DAPI. Images were acquired at 63X magnification using a Zeiss Axio Imager M2 microscope. Insets show magnified views of the indicated regions. (The Control and Act D treated immunofluorescence images shown here are the same representative images presented in Figure 1D).

The translocation of HuR between the nucleus and cytoplasm is mediated via the HuR Nucleocytoplasmic Shuttling (HNS) sequence. The HNS is a 33-amino acid domain (residues 205–237) located in the hinge region between RNA recognition motifs 2 (RRM2) and 3 (RRM3) of the HuR protein (40). To further delineate the role of the cytoplasmic translocation of HuR, we investigated the consequences of the removal of the HNS of HuR (HuR-ΔHNS) on its cytoplasmic localization in comparison with the full-length protein (HuR-FL) in the presence of Act D or vehicle. Both proteins were tagged with GFP. HuR-FL showed cytoplasmic accumulation with Act D treatment, as expected; however, HuR-ΔHNS was distributed both to the nucleus and the cytoplasm regardless of Act D treatment (Figure 2C, Figure S6A, B). This suggests that although the HNS region is important for the subcellular localization of HuR, it did not mediate the cytosolic localization in the presence of Act D.

We next evaluated whether we could inhibit the cytoplasmic shuttling of HuR upon Act D treatment. For this, we inhibited nuclear export with Leptomycin B (LMB). Although LMB treatment led to the nuclear accumulation of the positive control p53 (Figure S7A), we did not observe any effect on the cytoplasmic localization of HuR (Figure 2D, Figure S7B, C). Immunofluorescence images also revealed the cytoplasmic translocation of HuR in the presence of both Act D and LMB. This suggested that rather than enhancing the cytoplasmic shuttling of HuR, Act D may have enhanced the sequestration of HuR in the cytoplasm. Of note, we observed the HuR protein in bright, punctate/granular structures in the cytoplasm (Figure 2C&D, arrows in insets) hinting at the possibility of clustering of HuR in the cytosol.

### 3.3 Identification of Proteins in Close Proximity to HuR During Act D Induced Stress Implications of Granule Formation

We next queried whether the existing cytosolic mRNAs were necessary for the formation of the granules. For this, we permeabilized the cells with Tween 20 and treated the cells with RNase A as described in (41) or Act D (Figure S8). RNAse A treatment retained cytoplasmic HuR at levels comparable to that of Act D treatment alone, suggesting that neither newly synthesized, nor existing RNAs were necessary for generating the granular structures. This suggested that protein-protein interactions might have more important contribution to HuR-associated granule assembly, without excluding a role for RNA.

Therefore, we next evaluated whether HuR may be sequestered with other proteins in the cytoplasm upon Act D treatment. To this end, we carried out a TurboID-based proximity biotinylation assay to identify proteins in close proximity to HuR. Variables such as amount and duration of incubation with biotin were optimized (Figure S9A, B). We first confirmed that the TurboID tagged HuR protein also showed cytoplasmic translocation when treated with Act D (Figure S9C). The LC-MS/MS assay carried out after pulldown of the proteins with streptavidin, led to the identification of 52 proteins that were specifically biotinylated by HuR-TurboID and therefore were in the close vicinity to HuR only upon Act D treatment (Figure 3A, Figure S10A for the complete list of proteins).

**Figure 3.**
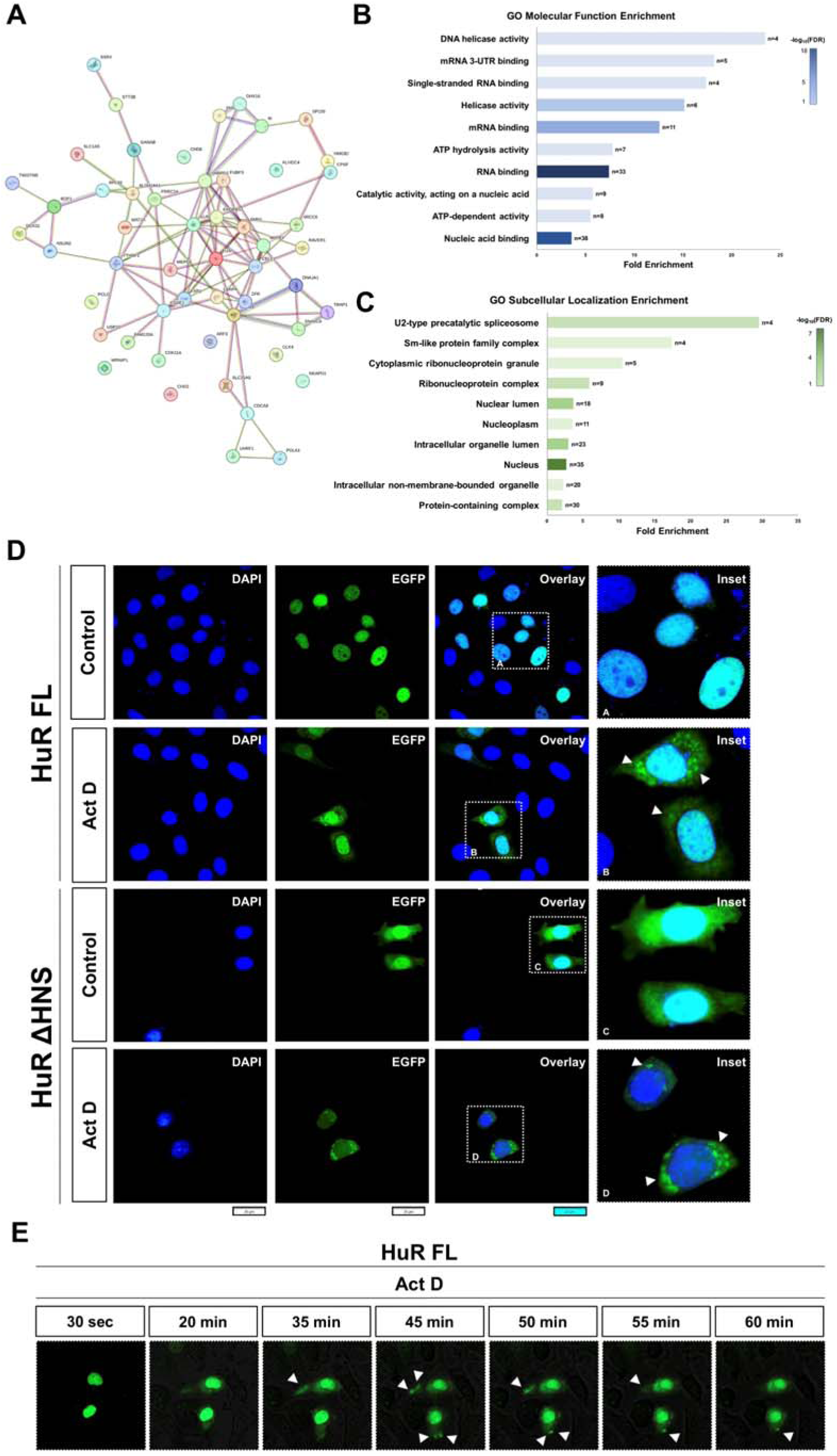
HuR Phase Separates into Dynamic Cytoplasmic Condensates Possibly with Other RNA-Binding Proteins Based on the HuR Proximity Interactome. (A) A total of 52 proteins biotinylated by HuR-TurboID in the presence of Actinomycin D (Act D) were identified and visualized as an interaction network. Data from two independent biological replicates were analyzed. Gene ontology (GO) enrichment analysis of the identified proteins was performed using the STRING database. Enrichment results for molecular function and subcellular localization are presented in (B) and (C), respectively, showing the top 10 enriched terms ranked by fold enrichment. (D) HeLa cells were transfected with pEGFP-c1-HuR FL or pEGFP-c1-ΔHNS constructs for 24 h. Cells were subsequently treated with 10 μg/mL Actinomycin D (Act D) for 3 h or left untreated as controls. GFP fluorescence was imaged using an Olympus BX43 microscope. (E) HeLa cells were transfected with pEGFP-c1-HuR Full-Length (EGFP-HuR) for 48 h. Transfected cells were treated with 10 μg/mL Actinomycin D (Act D) for 1 h and subjected to live-cell imaging during the treatment using a Zeiss Axio Imager M2 microscope, with images acquired every 30 seconds. Representative time-lapse images are shown.

The gene ontology enrichment profiling of the 52 proteins according to their molecular function (Figure 3B, Figure S10C) and biological processes (Figure S10B) indicated a strong enrichment of proteins with RNA-related functions, such as RNA-binding, processing, and stabilization, together with regulatory processes such as RNA splicing and post-transcriptional gene regulation. Analysis of the subcellular localization (Figure 3C, Figure S10E) and cellular component (Figure S10D) showed that the HuR-proximal proteins were also associated with non-membrane bound complexes such as spliceosome, stress granules, and ribonucleoprotein complexes. These data strongly indicated that HuR could interact with, or reside in close proximity to other RBPs, potentially forming granular condensates/aggregates in response to the stress of transcription inhibition (28, 29). To evaluate the formation of granular structures, we transfected HeLa cells with the GFP-tagged HuR-FL or ΔHNS constructs and treated the cells with Act D for 3 h. We observed that both full length and truncated HuR proteins formed bright, puncta-like structures in the cytoplasm only when treated with Act D (Figure 3D). Moreover, live cell imaging using GFP-tagged HuR FL (Figure 3E) indicated that granular structures were formed within 45 mins of treatment of HeLa cells with Act D, which later fused with each other, exhibiting the typical temporal behavior of cellular aggregates (Supplementary Video 1) (23). Supporting this, we observed that short-term treatment (15 min, 30 min, 1h and 2h) of both cell lines with Act D increased the cytoplasmic levels of HuR (Figure S11).

Since many RBPs are known to form granules in the cytoplasm or nucleus (42, 43), we reasoned that Act D treatment may lead to the cytosolic sequestration of multiple RBPs. Indeed, we observed that in addition to HuR, the RBPs CUGBP1, CUGBP2, TIAR, AUF1 and FUS also showed increased cytoplasmic levels and/or decreased nuclear levels upon Act D treatment (Figure S12 A&B).

### 3.5 Probabilistic modeling for the granule forming ability of RBPs in the presence of Act D

We next evaluated which proteins in close proximity to HuR could participate in the formation of granular structures. For this, we used the STRING analysis outputs of the TurboID data and the available literature to categorize the HuR-interacting proteins according to the following criteria: direct interaction with HuR, known association with granules, the presence of IDRs in their protein structure and having an isoelectric point (pI) higher than physiological pH (carry a net positive charge) (Figure S13). We determined the pI of the proteins since condensates were recently shown to function as spatially compartmentalized buffers in which charged proteins, RNA and other components could contribute towards the acid-base equilibrium (44).

We next developed a probabilistic framework integrating experimental data with Monte Carlo simulation and Markov state-transition modeling to identify the proteins most closely associated with HuR and estimate their probabilities of stable granule formation (Figure S14). Average Assembly Scores calculated across Monte Carlo simulations differed among the HuR-associated proteins (Figure S15A). These scores were subsequently incorporated into the Markov state-transition model to estimate the probability of stable granule formation (Figure S15B). FUS exhibited the highest Assembly Score, followed by CELF1, KHDRBS1, SNRPD2, YTHDF2 and FUBP3, whereas PRRC2A, CPSF7, and ZFR consistently showed lower scores. Stable granule probabilities closely reflected the Assembly Score rankings, with FUS and CELF1 displaying probabilities approaching 1, whereas PRRC2A, CPSF7, and ZFR exhibited markedly lower probabilities.

We next carried out a sensitivity analysis to quantify the relative contribution of normalized isoelectric point, condensate localization, PPI score and IDR annotation to the predicted probability of stable granule formation (Figure S15C). PPI showed the strongest positive association with stable granule formation, whereas condensate localization showed a weaker but consistently positive contribution. In contrast, IDR annotation and normalized pI demonstrated negative correlations within the current model. Given the limited number of proteins and the dependency introduced by Dirichlet-based weight sampling, these negative correlations should be interpreted cautiously.

The relationship between experimental features and computational predictions was visualized via two complementary representations (Figure 4). The bubble plot demonstrated a strong positive relationship between the mean Assembly Score and Stable Granule probability, with bubble size representing Assembly Score magnitude and color indicating the strength of interaction in the protein network (Figure 4A). FUS and CELF1 occupied the upper-right region of the plot, consistent with their high Assembly Scores and Stable Granule probabilities. The heatmap (Figure 4B) summarized the experimental variables together with the predicted Assembly Scores and Stable Granule probabilities. Proteins ranked highly by the computational framework consistently displayed stronger interaction scores and more favorable feature profiles than lower-ranked proteins, highlighting good consistency between experimental observations and probabilistic predictions.

**Figure 4.**
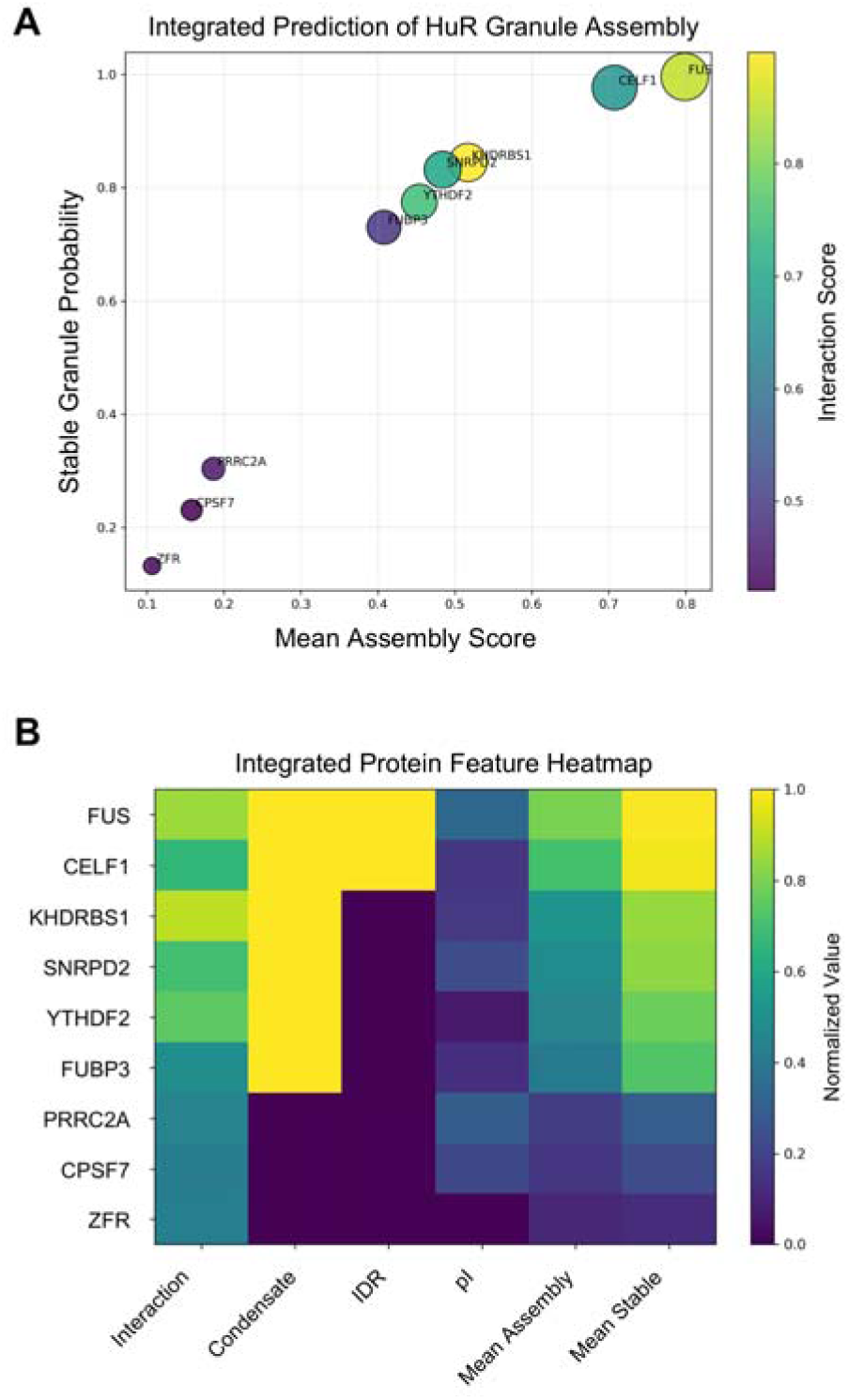
Integrated visualization of experimental features and model predictions. (A) Bubble plot showed the relationship between mean Assembly Score and Stable Granule probability, with bubble size proportional to Assembly Score and color representing PPI score. (B) Heatmap summarized normalized experimental features together with predicted Assembly Scores and Stable Granule probabilities.

### 3.4 HuR-FUS Interaction in the Presence of Act D

Based on the probabilistic modeling, we selected FUS as the candidate protein to validate our data as the protein has been widely reported to localize to multiple distinct condensates both in the nucleus and cytoplasm (45–48). FUS is predominantly a nuclear protein (49); however, we observed that in the presence of Act D, the FUS protein could also translocate to the cytoplasm (Figure 5A). To understand whether HuR and FUS could physically interact, we carried out protein-protein-docking and molecular dynamics simulations. The docking model supported the formation of a plausible interaction interface between the two proteins (Figure 5B). Moreover, when we incorporated two Act D molecules into the HuR-FUS complex (Figure S16), the mean square displacement (MSD) analysis of the HuR-FUS interaction showed that structural fluctuations were limited throughout the simulation period (Figure 5C). This suggested that the HuR-FUS interaction was dynamically stable. Comparative analysis of MSD trajectories of the interaction between HuR-FUS and HUR-FUS-2Act D systems revealed similar dynamic behaviors (Figure 5C) suggesting that the incorporation of Act D did not alter the structural stability of the system in the simulation. Importantly, the Act D incorporated HuR-FUS complex exhibited a more favorable energy profile compared to HuR-FUS interaction alone, supporting enhanced interactions between these RBPs in the presence of Act D.

**Figure 5.**
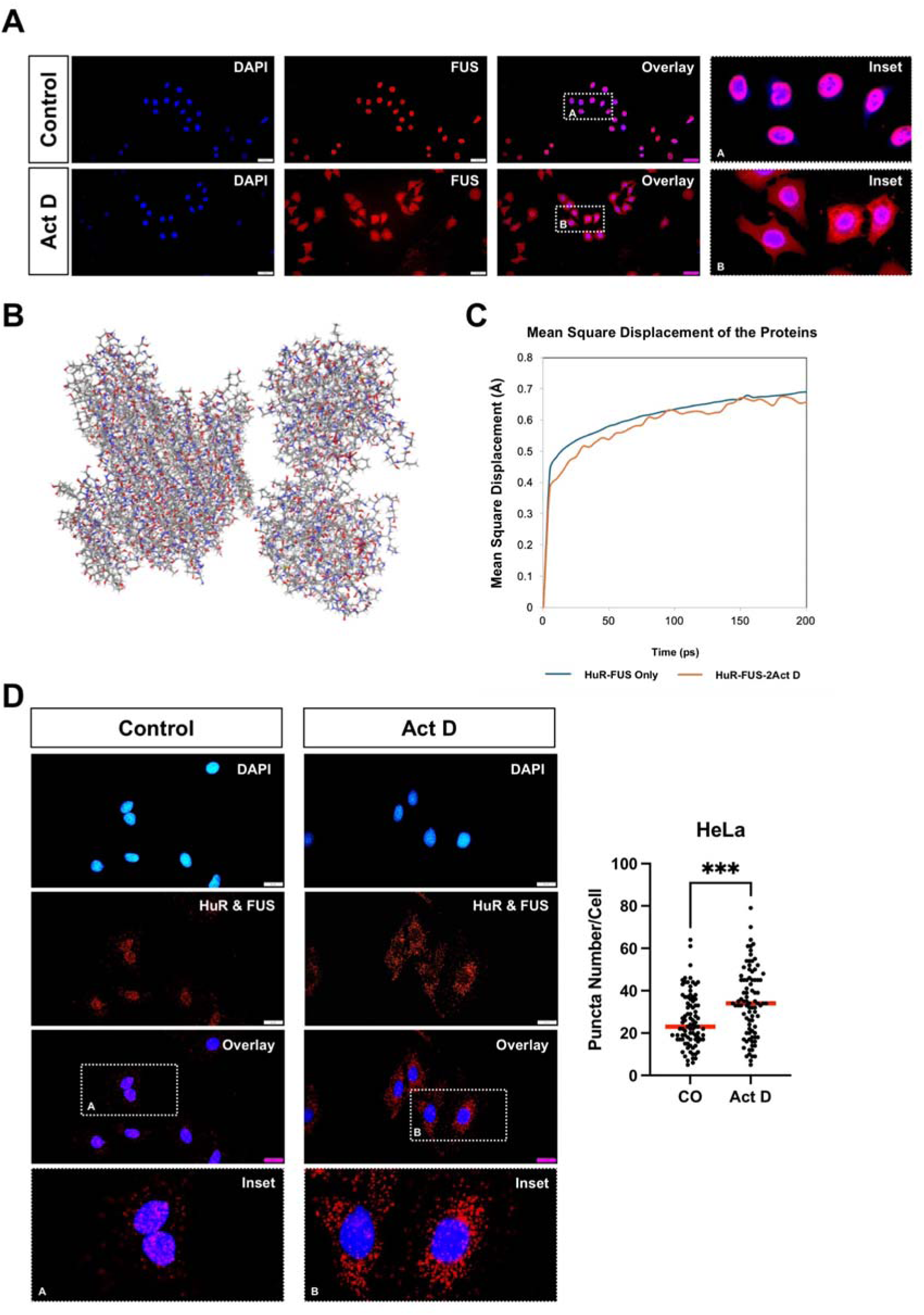
Validation of interaction between HuR and FUS in the presence of Act. **D.** (A) Immunofluorescence staining of HeLa cells demonstrates the translocation of FUS to the cytoplasm following Actinomycin D (Act D) treatment. Scale bar, 50 μm. (B) Protein-protein docking analysis was carried out to model the interaction between HuR and FUS. The model showed a binding energy of -2.443 kcal/mol, suggesting a potential interaction between the two proteins. (C) Molecular dynamic simulations were performed by utilizing the COMPASS II force field under the NPT ensemble conditions. The dynamic behavior and stability of the system were determined via the analysis of mean square displacement (MSD). A more favorable energy profile was observed when Act D was incorporated into the model (D) HeLa cells were treated with Act D or vehicle for 3 h, fixed, and incubated with antibodies against HuR and FUS. Cells were subsequently processed for proximity ligation assay (PLA). Two independent biological replicates were analyzed. Puncta quantification was performed on a total of 88 cells per experimental group. Statistical significance was determined using an unpaired t-test (***p < 0.001). Insets show magnified views of the indicated regions. The images were acquired with an Olympus BX43 microscope.

Next, we used proximity ligation assay (PLA) to validate the potential interaction between native HuR and FUS proteins. The assay demonstrated significantly increased interactions when cells were treated with Act D relative to vehicle-treated controls (Figure 5D). Cells incubated individually with the HuR or FUS antibodies showed very low number of red puncta supporting the specificity of the proximity labelling of HuR and FUS (Figure S17). Taken together, these data suggested that treatment with Act D induced the cytoplasmic translocation of several different RBPs including HuR and FUS, and that the latter two proteins were indeed in close proximity to each other.

### 3.4 Selective Dissolution of Granular HuR Structures by Hypotonic Shock

Building on our observation of granular structures with GFP-tagged and endogenous HuR in the cytoplasm and its proximity to FUS and other RBPs, we next tested whether the interaction relied on the presence of Act D, i.e., whether removal of Act D could lead to a reversal in the cytosolic accumulation of RBPs (50, 51). We observed no change in the cytoplasmic levels of HuR and FUS for up to 16h after the washout of Act D, suggesting the presence of robust interactions (Figure 6A, B).

**Figure 6.**
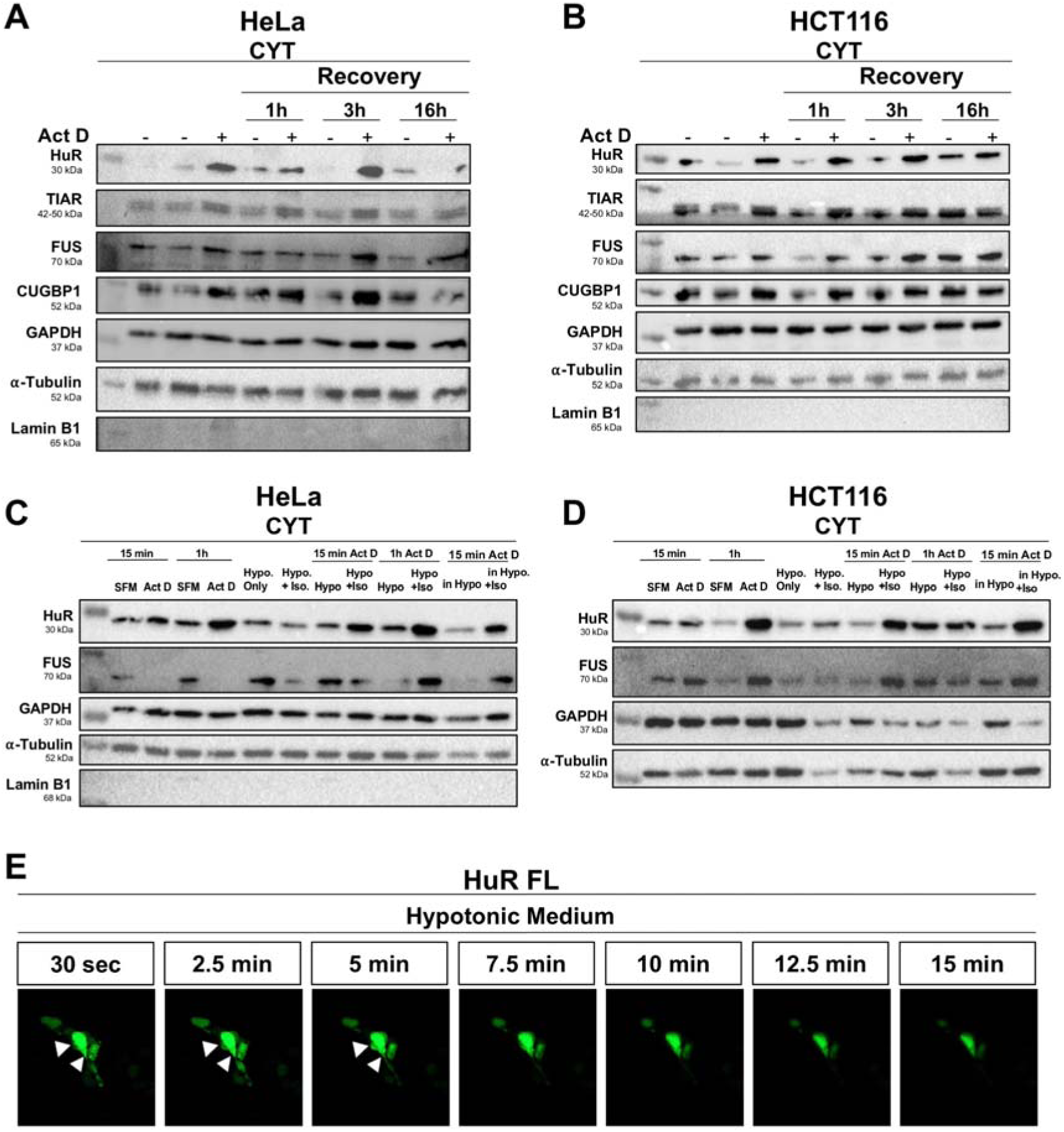
(A-B) HeLa (A) and HCT116 (B) cells were treated with Actinomycin D (Act D) for 3 h, followed by recovery in complete medium for 1 h, 3 h, or 16 h. Cytoplasmic (CYT) protein levels of HuR, FUS, TIAR, and CUGBP1 were analyzed. GAPDH and α-tubulin were used as cytoplasmic loading controls, while Lamin B1 served as a nuclear loading control. (C-D) HeLa (C) and HCT116 (D) cells were incubated in hypotonic medium (Hypo) for 15 min following Act D treatment for either 15 min or 1 h. To detect reversibility, cells were returned to isotonic medium (Iso) for 1 h after hypotonic treatment (Hypo+Iso). As an additional control, Act D treatment was also performed under hypotonic conditions, followed by incubation in isotonic medium for 1 h. The durations of Act D treatments are indicated above the corresponding labels. Cytoplasmic levels of HuR and FUS were analyzed, with α-tubulin used as a loading control for cytoplasmic fractions. (Representative blot shown for at least two biological replicates) (E) HeLa cells were transfected with pEGFP-c1-HuR Full-Length (EGFP-HuR) for 48 h. After transfection, cells were treated with 10 μg/mL Actinomycin D (Act D) for 1 h, followed by incubation in hypotonic medium for 15 min. Live-cell imaging was performed during the hypotonic treatment using a Zeiss Axio Imager M2 microscope, with images captured every 30 seconds. Representative time-lapse images are shown.

Incubation of cells in a hypotonic environment is known to enhance cellular swelling through osmosis; this process is also widely known to disrupt condensates (52). We observed that incubation of cells in a hypotonic medium for only 15 min after Act D treatment resulted in a decrease in the cytoplasmic levels of both HuR and FUS (Figure 6C, D). Furthermore, restoration of the isotonic medium in the presence of Act D could also restore the HuR and FUS aggregates in the cytoplasm (Figure 6C, D). A live cell imaging assay revealed that incubation of Act D treated cells in the hypotonic medium not only disrupted the granules but also decreased the cytoplasmic levels of HuR (Figure 6E, Supplementary Video 2). These findings suggest that upon Act D treatment, HuR and other RBPs can form granules in the cytosol that are relatively dynamic but remained in the cytoplasm long after the removal of Act D.

1,6-hexanediol (HDO) is an alcohol that is widely used to disrupt weak protein-protein interactions in condensates (53). We next tested whether the cytoplasmic accumulation of the RBPs could be reversed with HDO. We selected 2.5% HDO for 5 min as the optimized condition that did not affect cell viability or induce detectable morphological alterations (Figure S18A). Use of higher concentrations or longer durations led to extensive cell death. These conditions did not disrupt sodium arsenite-induced stress granule formation (Figure S18B); therefore, we were not able to conclude whether treatment with HDO could inhibit the cytoplasmic accumulation of FUS and HuR.

## 4 Discussion

Cells activate various signaling pathways in response to stress. The current study was designed to understand why RBPs in epithelial cells localize to the cytosol upon global transcriptional inhibition with Act D. Evaluation of a publicly available RNA-seq dataset (GSE198178) (34) showed that a distinct population of mRNAs were upregulated upon treatment of HeLa cells with 5μM Act D (0h, 1h, 2h, 4h), despite global transcriptional inhibition. Gene ontology enrichment analysis revealed that Act D treatment could induce a robust ribosomal stress response characterized by the enrichment of genes in ribosome biogenesis, translation as well as RBPs reflecting a compensatory adaptation to impaired transcription (54).

We focused on HuR (*ELAVL1*), an RBP that plays an important role in increasing mRNA stability and protein translation (55, 56). Our initial observations demonstrated that the predominantly nuclear HuR protein undergoes rapid and robust cytoplasmic translocation following Act D treatment. To understand the mechanism, we first focused on DNA damage response (DDR) pathways, since Act D can intercalate into DNA and activate DDR. Additionally, Chk2, a critical kinase activated in DDR pathways (57), has been reported to promote cytosolic translocation of HuR and stabilization of its target mRNAs (58). However, treatment of cells with multiple DNA damaging agents did not lead to the cytosolic translocation of HuR, suggesting that the phenomenon was not a response to DNA damage.

We next investigated the involvement of nuclear export pathways in the cytosolic translocation of HuR. HuR has been reported to undergo CRM1-dependent nuclear export (59, 60), which can be inhibited by leptomycin B (LMB) (61). Surprisingly, LMB treatment could not inhibit the Act D-induced cytoplasmic localization of HuR, suggesting that its export may occur via CRM1-independent mechanisms (62, 63). The HuR nucleocytoplasmic shuttling (HNS) sequence mediates the bidirectional transport of the protein (10). Deletion of the HNS domain (ΔHNS) resulted in equal distribution of HuR in the nucleus and cytoplasm under basal conditions, while full-length HuR remained predominantly nuclear. Both proteins translocated to the cytosol upon Act D treatment, suggesting that the HNS domain was not critical for Act D mediated cytosolic translocation.

HuR was observed in puncta like aggregates in the cytosol, which increased in size in a temporal manner. Since proteins are known to be transiently sequestered into stress-induced biomolecular condensates (42, 64), we reasoned that rather than increasing the cytoplasmic translocation, Act D treatment may lead to the sequestration of HuR in the cytosol. The HuR aggregates were not likely to be condensates since Act D treatment is known to decrease the formation of P-bodies (65, 66) and stress granules (67) by decreasing the availability of RNA substrates that are required for condensate assembly. Supporting this, we did not observe any G3BP1 positive stress granules in cells treated with Act D for 1h (Figure S19); additionally, the treatment of the cells with RNAse A did not impede the cytosolic localization of the RBPs.

The IDR prediction tools PONDR and IUPRED3 revealed the presence of structured regions in the HuR protein that are connected by flexible disordered regions that most likely facilitate protein-protein interactions (Figure S20). Moreover, we identified several different RBPs in close proximity to HuR only when the cells were treated with Act D. Probabilistic modeling indicated the proteins FUS and CELF1 to have the highest probability of being in close proximity to HuR. FUS which has a well-established role in forming aggregates (68–71) and was observed to translocate to the cytoplasm upon treatment with Act D. More importantly, both MD simulations and a proximity ligation assay could confirm the interaction between the two proteins only when Act D was present. Taken together, our data strongly pointed towards the rapid formation of RBP aggregates in cells treated with Act D.

Several studies have shown that removal of the stress can lead to rapid disassembly of condensates (50, 51). On the other hand, the stress granule protein TDP-43 was shown to persist in the form of aggregates in the cytoplasm even after the removal of the stress (72). We observed by western blot that the elevated cytoplasmic levels of several condensate-associated RBPs such as HuR, FUS, CUGBP1 and TIAR (73–75) remained even after stress removal. Time lapse imaging indicated that the aggregates were able to increase and then decrease in size, suggesting the presence of a level of fluidity. Of note, treatment of the cells with the detergent Tween 20 alone, which can generate micron sized pores in the plasma membrane (41), resulted in the retention of the RBPs in the nucleus and/or loss from the cytosol. However, when we co-treated the cells with Tween 20 and Act D, we again observed the cytosolic translocation and retention of the RBPs in the cytosol (Figure. S8). This suggested that the proteins aggregates were larger than the size of the pores generated with Tween 20.

The aggregation and disaggregation of mammalian proteins can be mediated by heat shock proteins and chaperones; these proteins sequester and stabilize aggregation-prone proteins in response to stress and then allow their refolding via the HSP70 family of chaperones when the stress is relieved (76). We observed a number of heat shock proteins (DNAJC9, DNAJA1 and HSPA9) in close proximity to HuR only in cells treated with Act D. Several of the heat shock protein genes also showed a temporal increase in expression (GSE 198178) in HeLa cells treated with Act D (data not shown). This suggests that the RBP aggregates underwent a relatively fluid aggregation and disaggregation, all the time staying within the cytoplasm.

Hypotonic shock rapidly increases cellular volume and can disassemble condensates (77). Further confirmation of the fluidity of the aggregates was shown when a rapid decrease in the cytoplasmic levels of both HuR and FUS was observed with hypotonic shock. Moreover, the RBPs were relocated to the cytosol and underwent aggregation as soon as the cells were incubated in isotonic medium.

Collectively, our findings suggest that HuR undergoes cytoplasmic sequestration in the presence of Act D and associates with FUS and additional RBPs to form stress-induced aggregates. Future studies will indicate whether these aggregated proteins are functionally active or not.

## Acknowledgements

The authors would like to acknowledge Çağrı Gündüz, Sümeyra Özmen and Leyla Atakishiyeva for technical assistance and Busra Kırım for performing the LC-MS/MS analyses. Dr Elif Erson is acknowledged for helpful discussions. The research was partly supported by TÜSEB (Grant no: 38044) and ODTÜ BAP (GAP-108-2023-11313) to S.B. and TÜBİTAK 122Z491 to AT.

## Abbreviations

α-AMA: α-Amanitin
Act D: Actinomycin D
Chk2: Cell cycle checkpoint kinase
CPT: Camptothecin
CUGBP1: CUG-binding protein 1
CYT: Cytoplasmic
FUS: Fused in Sarcoma
HNS: HuR nucleocytoplasmic shuttling sequence
HuR: Human antigen R
IDR: Intrinsically disordered region
LMB: Leptomycin B
NUC: Nuclear
PLA: Proximity ligation assay

