## Supplementary Data for "Actinomycin D Drives RNA-Binding Proteins into Dynamic Cytoplasmic Granules"

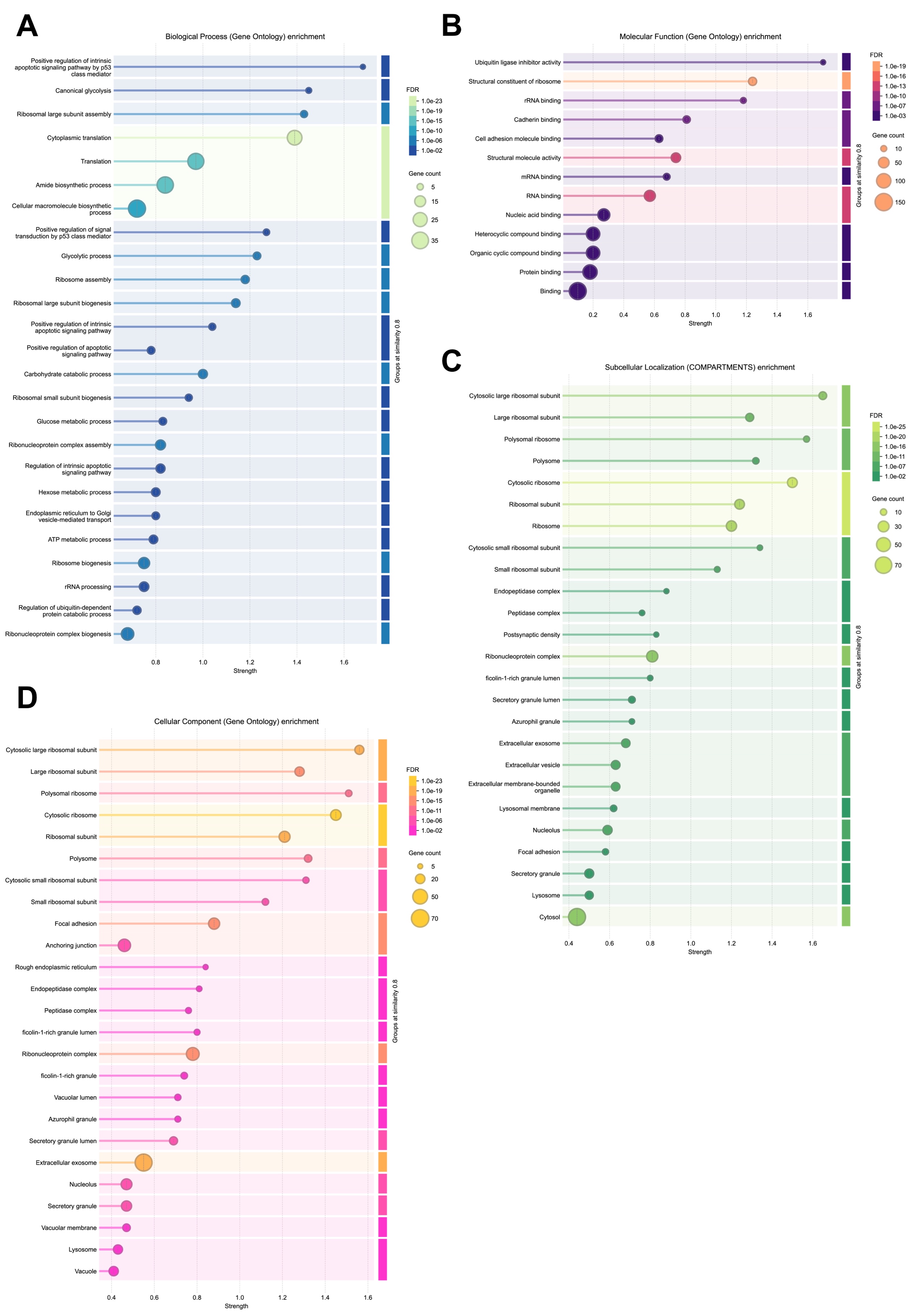

**Supplementary Figure 1: Gene ontology enrichment analysis of the differentially upregulated genes upon Act D treatment in the GSE198178 dataset was performed on STRING**. (A) GO biological process enrichment analysis revealed a significant enrichment of ribosome and translation related processes in upregulated genes in the presence of Act D (B) Molecular Function enrichment analysis of the upregulated genes showed enrichment of genes associated with ribosome and RNA binding activities. (C) Terms such as ribosome, cytoplasm, ribonucleoprotein complex were enriched in the Subcellular Localization analysis. (D) Cellular Component enrichment analysis revealed the enrichment of ribosome, polysome, as well as intracellular non-membrane bound organelle-related terms. (Only the top 25 terms are shown in the charts for clarity)

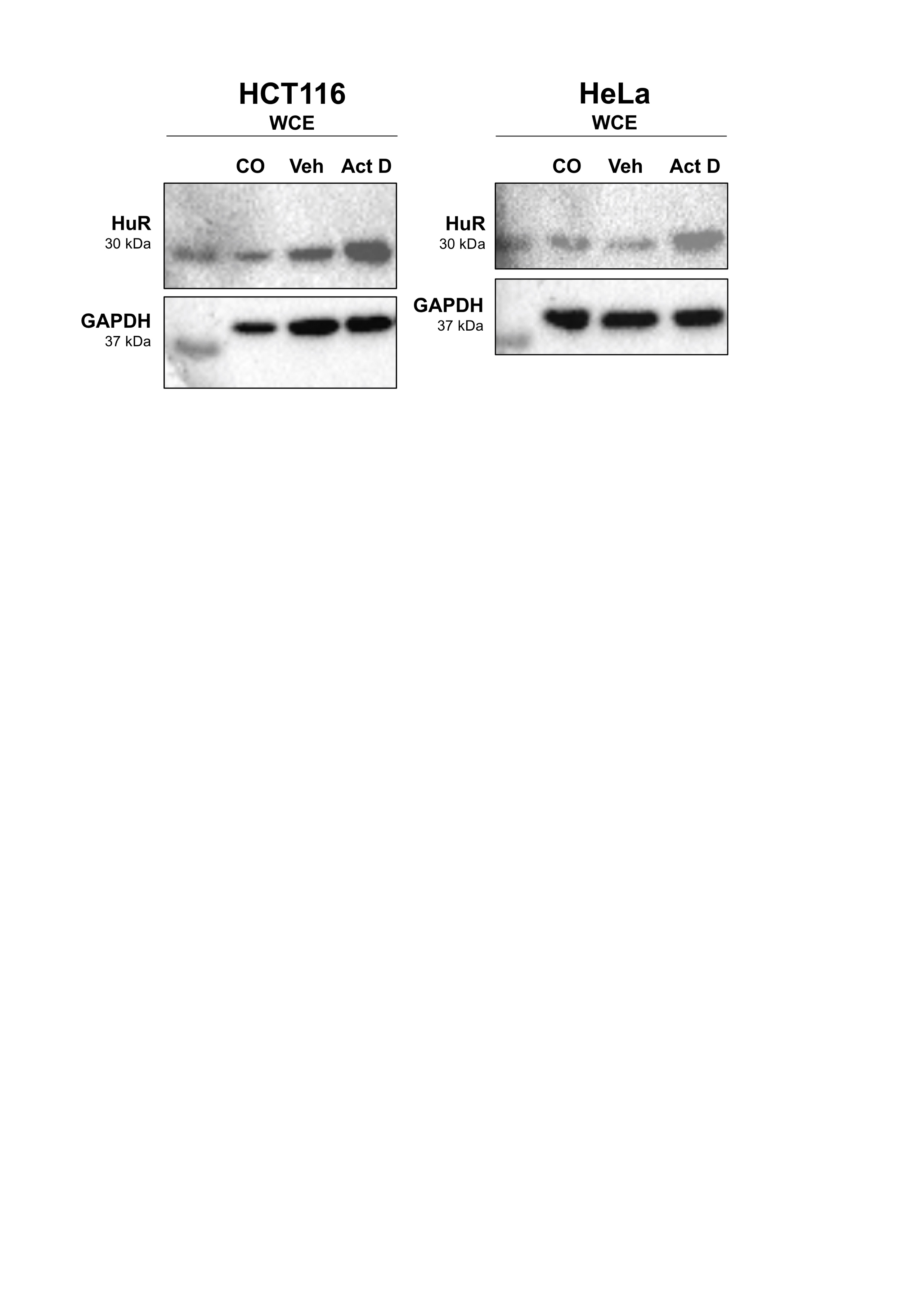

**Supplementary Figure 2: Total protein levels of HuR increased upon Act D treatment.** HCT116 and HeLa cells were treated with 10 μg/mL Act D for 3 h, followed by collection of whole cell extracts (WCE). Western blot analysis showed an increase in the total protein levels of HuR in the Act D treated cells compared to the untreated control (CO) and vehicle (Veh) treated groups.

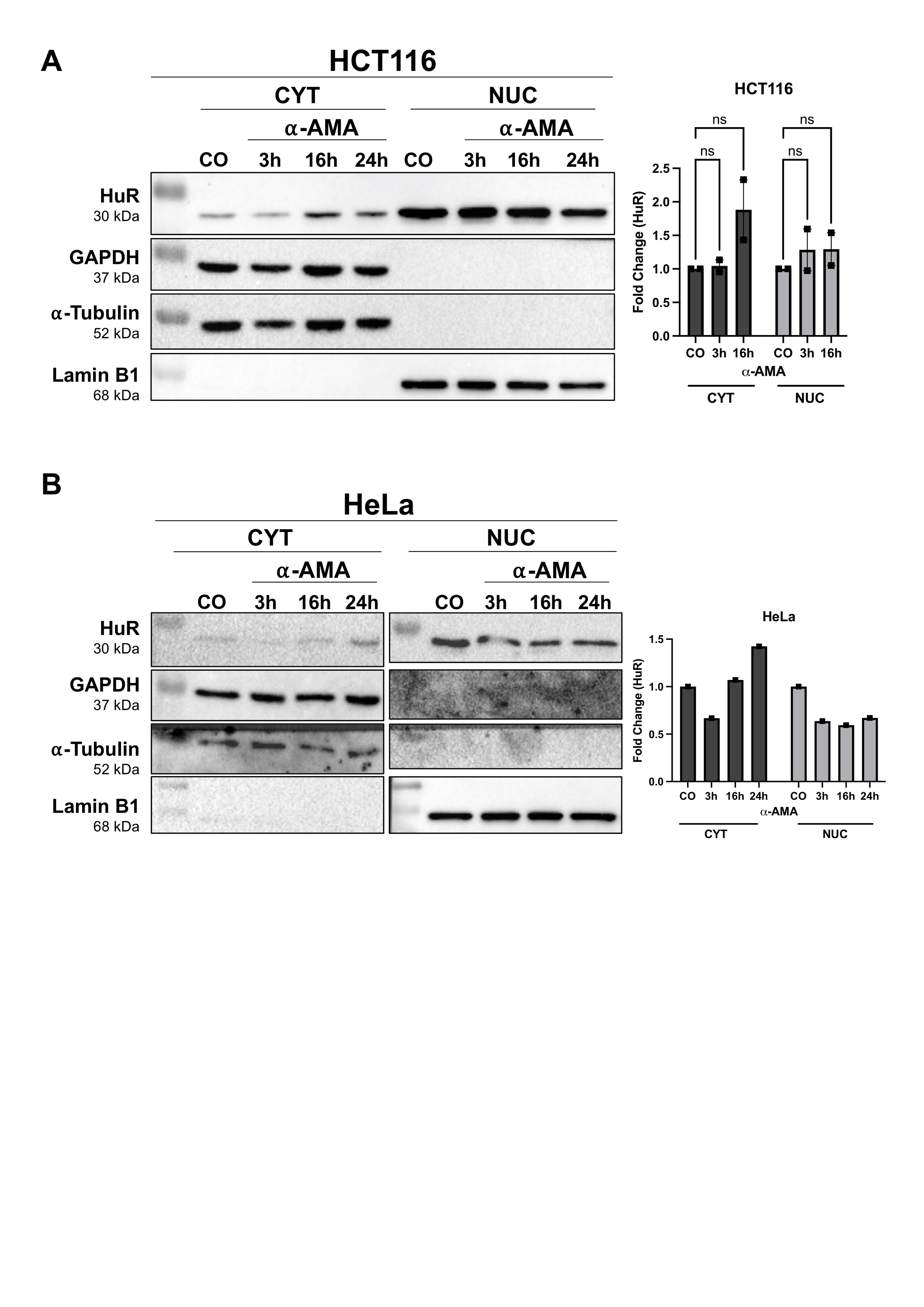
**Supplementary Figure 3. α-Amanitin (α-AMA) treatment required prolonged exposure to induce cytoplasmic localization of HuR.** HCT116 (A) and HeLa (B) cells were treated with 3 μg/mL α-AMA for 3, 16 and 24 h. Cytoplasmic (CYT) and nuclear (NUC) protein fractions were collected. GAPDH was used as loading control, α-Tubulin as a cytoplasmic marker and Lamin B1 as a nuclear marker. Densitometric quantifications were presented as bar graphs. Two independent replicates with HCT116 cells were carried out for statistical analysis. Fold changes were determined by normalization to untreated cells (CO). Statistical analyses were carried out by performing ANOVA with Tukey’s post hoc test. ns: not significant.

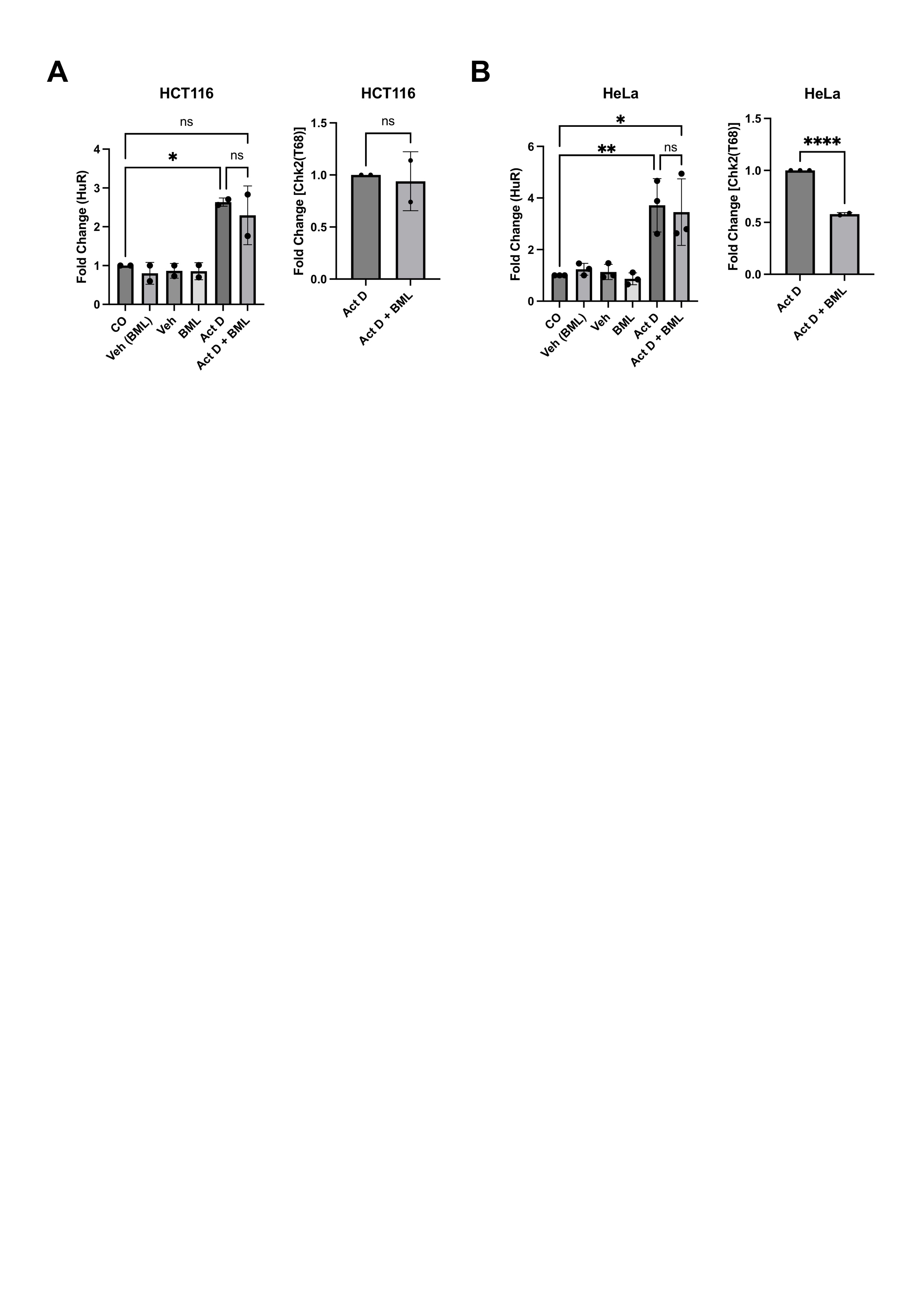

**Supplementary Figure 4. Chk2 inhibition did not alter the Act D-induced cytoplasmic localization of HuR.** Graphs represent the densitometric results of the experiments shown in Figure 2A&B and includes two biological replicates with HCT116 cells (A) and three biological replicates with HeLa cells (B). Statistical significance was determined by one-way ANOVA with Tukey’s post hoc test. *p<0.05, **p<0,005, ****p<0,0001, ns: not significant.

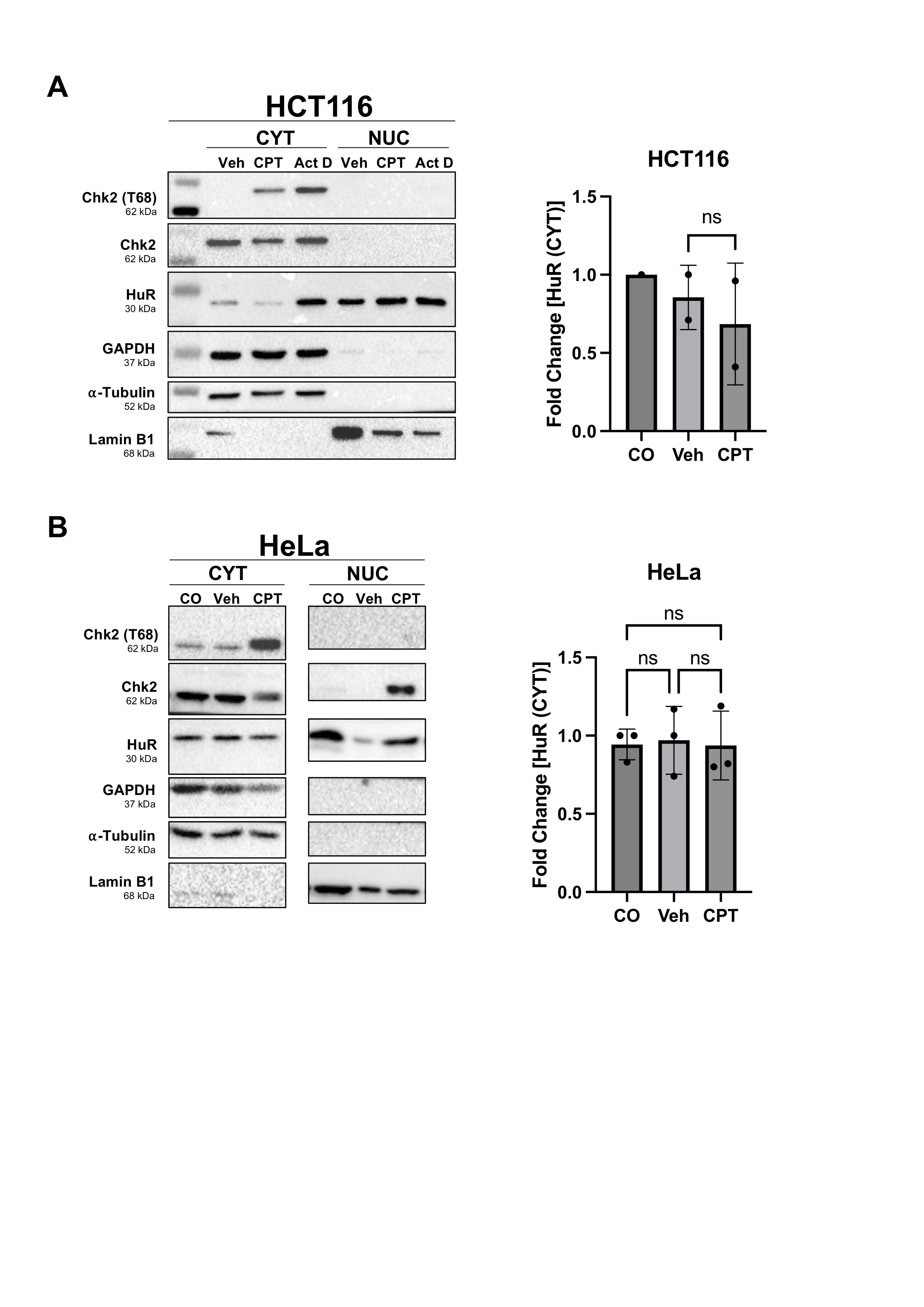

**Supplementary Figure 5. Camptothecin treatment did not affect the cytoplasmic localization of HuR.** HCT116 (A) and HeLa (B) cells were treated with 2 nM Camptothecin (CPT) for 24 h. Chk2 phosphorylation (T68) and HuR protein levels were analyzed in cytoplasmic (CYT) and nuclear (NUC) fractions. GAPDH and α-Tubulin were used as cytoplasmic while Lamin B1 was used as nuclear loading controls. Two independent biological replicates with HCT116 and three independent biological replicates with HeLa cells were carried out for statistical analysis. Bar graphs show the fold change in cytoplasmic levels of HuR. Statistical analysis was carried out via ANOVA with Tukey’s post hoc test. ns: not significant.

**Supplementary Figure 6. Removal of HNS region altered the subcellular distribution of HuR independently of Act D treatment.** HCT116 (A) and HeLa (B) cells were transfected with full length (HuR FL), ΔHNS or empty vector (EV, pEGFP-C1) constructs for 24 h. The cells were treated with 10 μg/mL Act D for 3 h. Cytoplasmic (CYT) and nuclear (NUC) fractions were isolated. The proteins were visualized by Western Blot. GAPDH, α-Tubulin and Lamin B1 were used as loading controls.

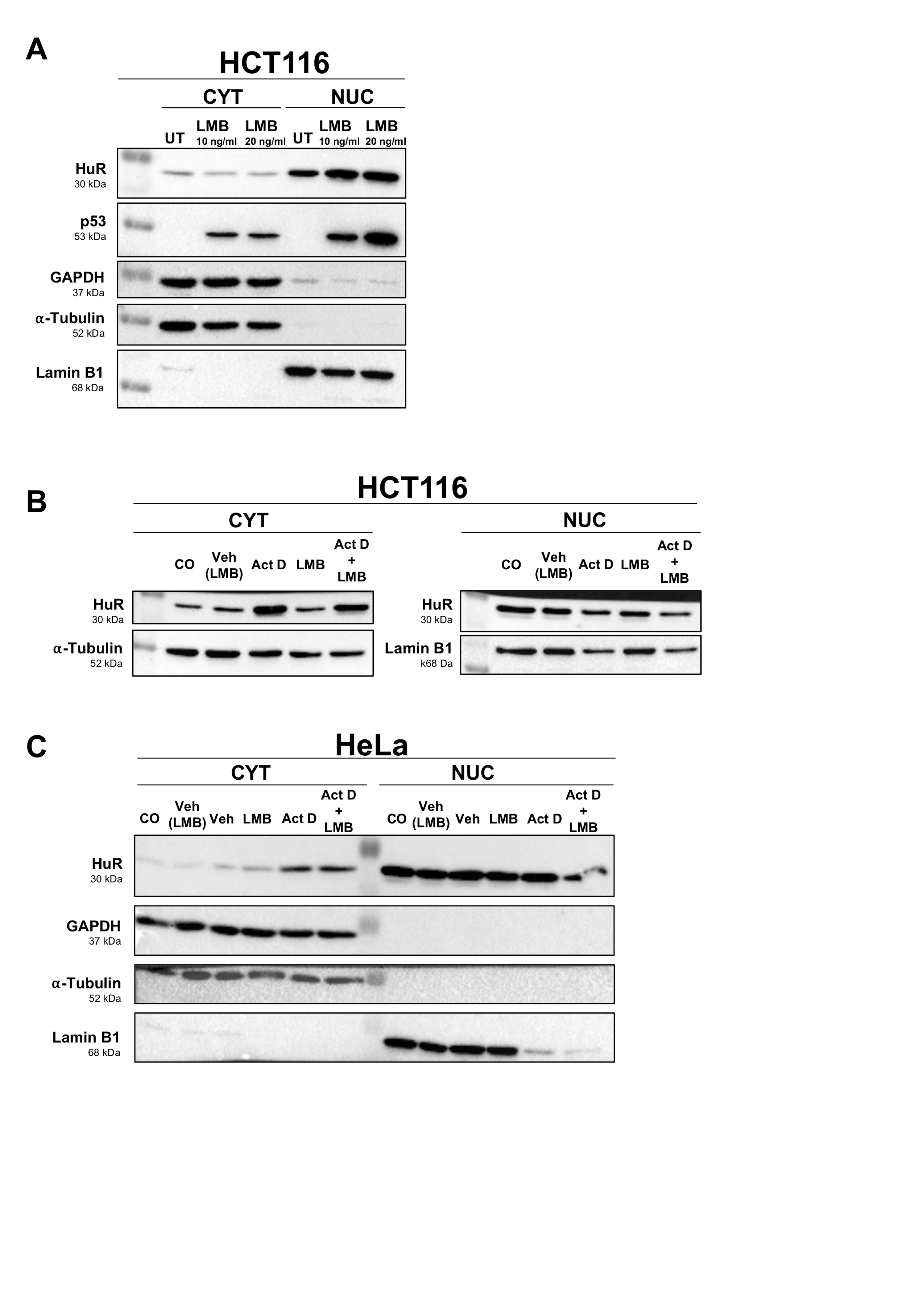

**Supplementary Figure 7. Leptomycin B (LMB) treatment did not reverse the effect of Act D on cytoplasmic HuR levels.** (A) Two different (10ng/mL & 20ng/mL) LMB concentrations were tested on HCT116 cells for 24 h to observe the effect of inhibition of nuclear export by LMB. Effect of Leptomycin B (LMB) was further evaluated with HCT116 (B) and HeLa (C) cells. The cells were treated with 10 ng/mL LMB for 24 h. For combined Act D and LMB treatment (Act D + LMB), cells were first incubated with LMB alone for 21 h, followed by co-treatment with Act D and LMB for an additional 3 h. Cytoplasmic (CYT) and nuclear (NUC) fractions were collected. Loading controls: GAPDH and α-Tubulin for CYT fractions, Lamin B1 for NUC fractions. (UT:Untreated, Veh: Vehicle).

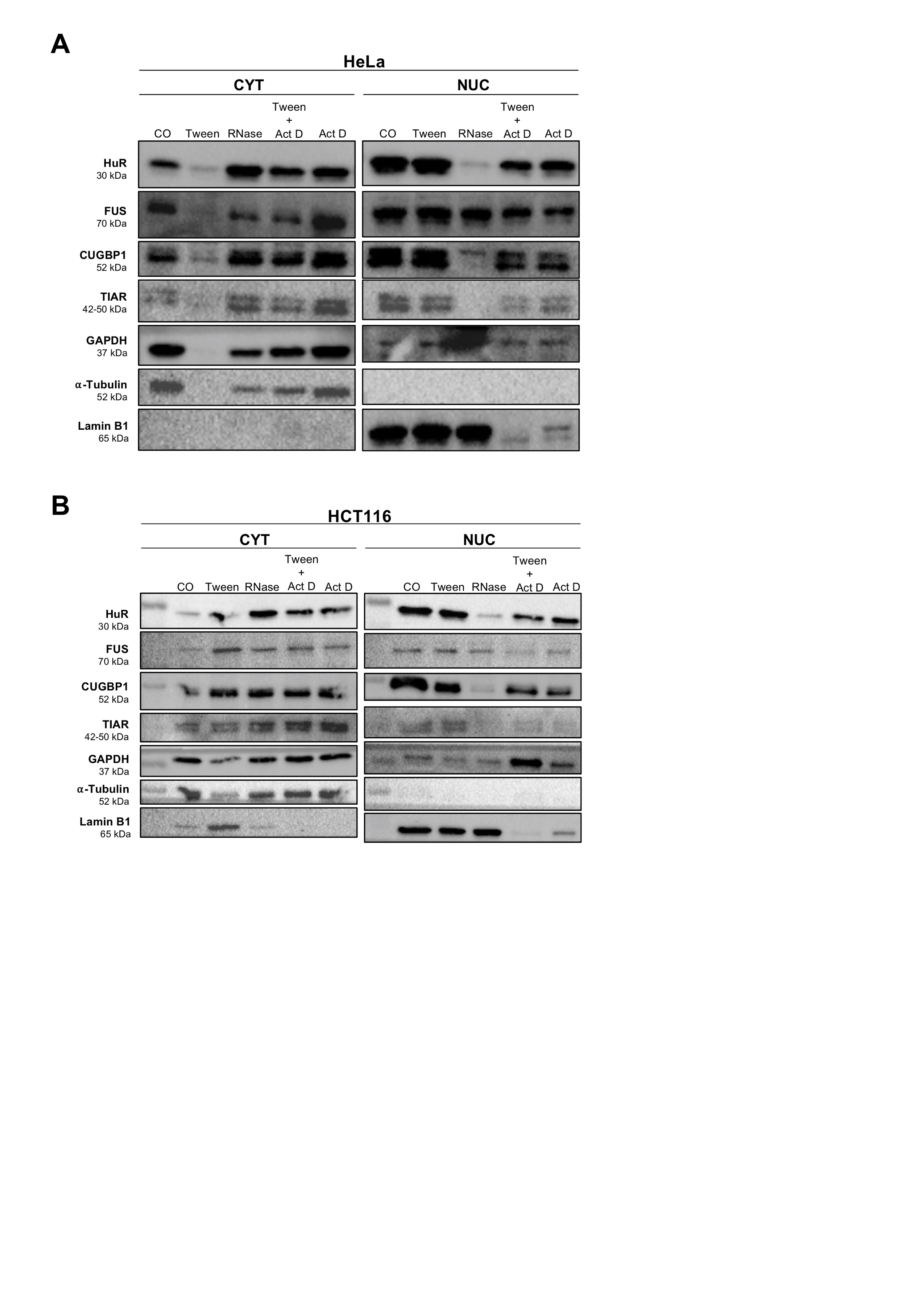

**Supplementary Figure 8. RNase A treatment increased the cytoplasmic levels of RBPs.** (A) HCT116 and (B) HeLa cells were permeabilized with 2% Tween 20 for 10 min. The cells were then incubated with 0.2 mg/mL RNase A for 30 min at RT on a shaker or with 10 μg/mL Act D at 37 °C for 1 h. Cytoplasmic (CYT) and Nuclear (NUC) protein fractions of the cells were collected after the indicated treatments. Loading controls: GAPDH and α-Tubulin for CYT fractions, Lamin B1 for NUC fractions.

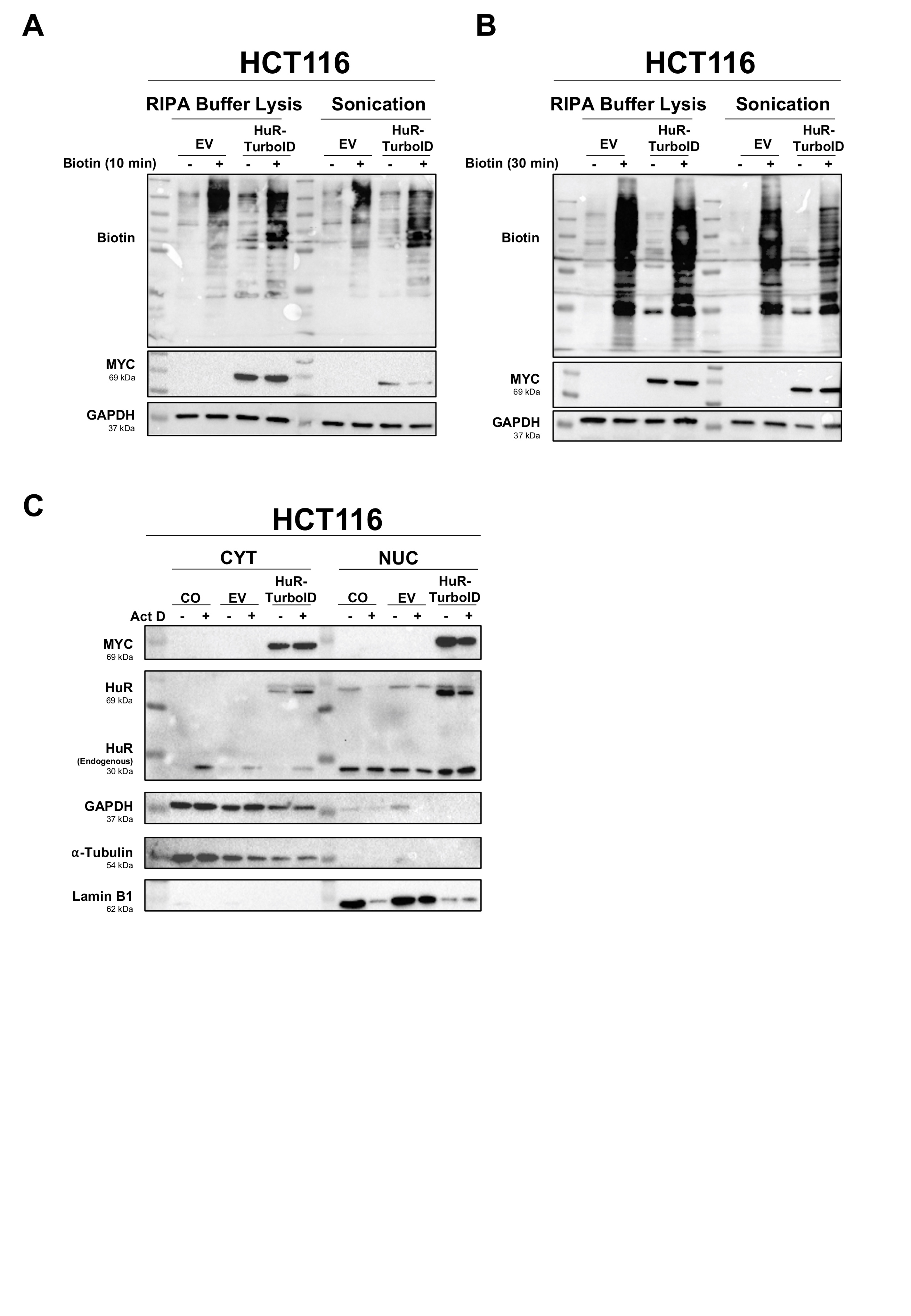

**Supplementary Figure 9. Optimization of experimental conditions for TurboID assay.** To determine the optimal biotin incubation duration, HCT116 cells that were transfected with EV or HuR-TurboID constructs were incubated with biotin for either 10 min (A) or 30 min (B). Next, the cells were lysed either via RIPA buffer or sonication. Protein biotinylation was analyzed by western blot probing with a biotin antibody. (C) HCT116 cells were transfected with pcDNA3.1 MYC-HuR-TurboID-HA (HuR-TurboID) or pcDNA3.1 TurboID-HA (EV) for 24 h. To test the functionality of the HuR-TurboID protein, cells were treated with Act D following transfection and separated into cytoplasmic and nuclear fractions. Protein levels of HuR-TurboID in subcellular fractions were determined using MYC or HuR antibodies. GAPDH and α-Tubulin were used as cytoplasmic controls and Lamin B1 as nuclear controls.

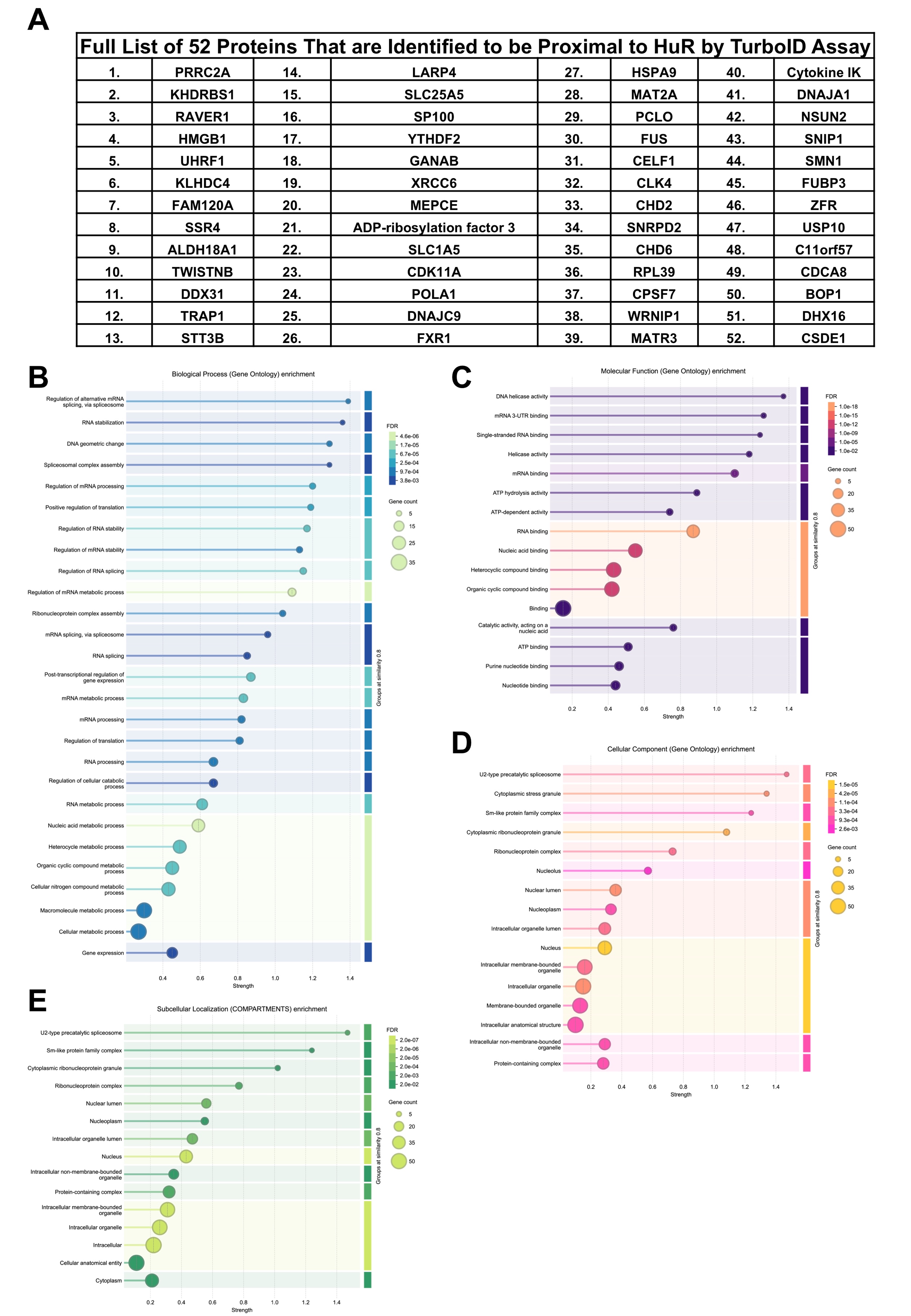

**Supplementary Figure 10. Proteins identified proximal to HuR were enriched in RNA regulation associated Gene Ontology (GO) terms.** (A) TurboID assay coupled with LC-MS/MS analysis identified 52 proteins biotinylated by HuR-TurboID in the presence of Act D. Results from two independent biological replicates were analyzed. GO enrichment analyses of the identified proteins were performed on STRING database for biological processes (B), molecular function (C), cellular component (D), and subcellular localization (E).

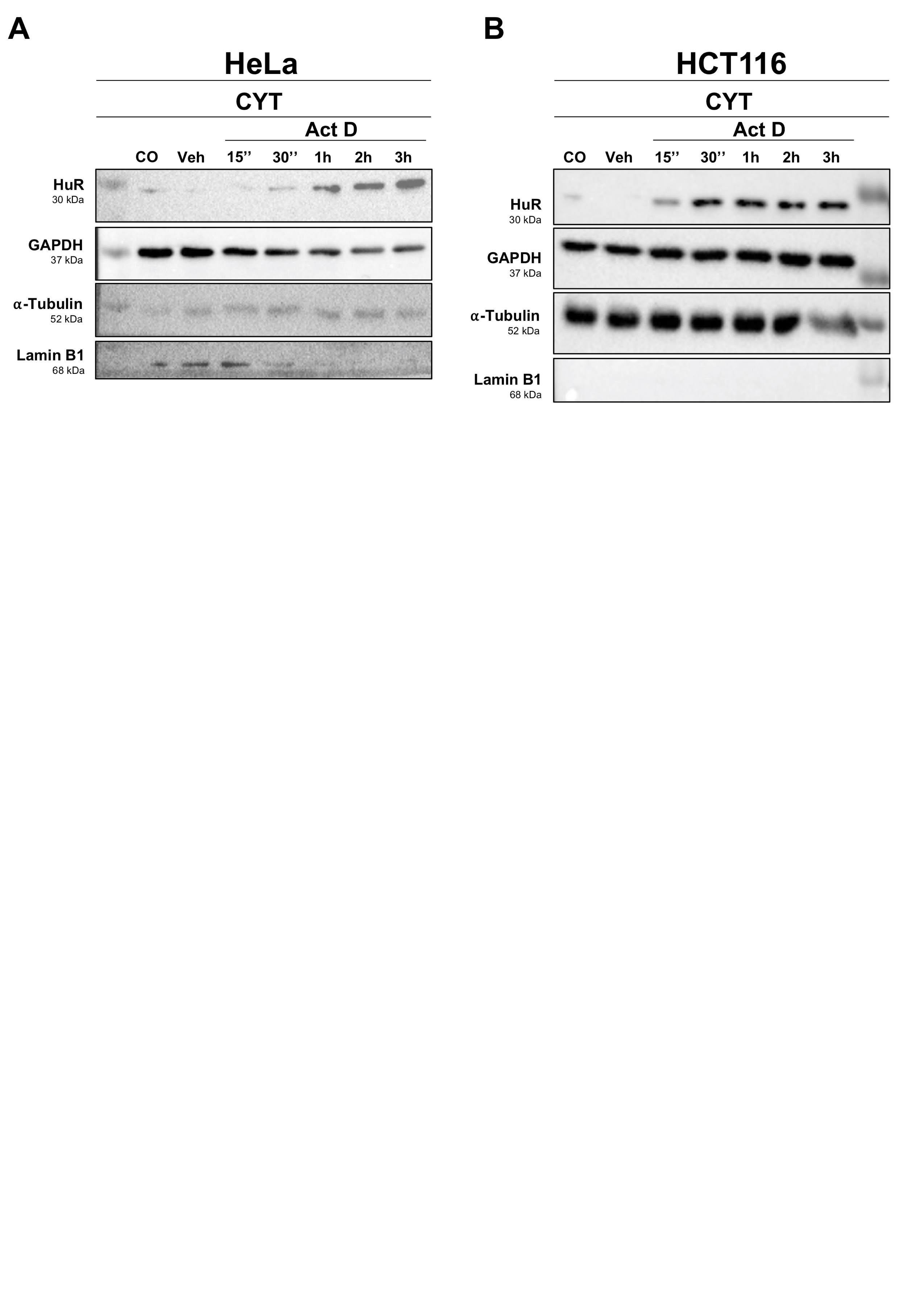

**Supplementary Figure 11. Short term treatment with Act D increased the cytoplasmic levels of HuR.** HeLa (A) and HCT116 cells (B) were treated with10 μg/mL Act D for 15 min, 30 min, 1 h, 2 h or 3 h. The subcellular protein fractions were collected. GAPDH and α-Tubulin were used as cytoplasmic controls and Lamin B1 as nuclear controls. CYT: cytoplasmic, CO: untreated cells, control, Veh: Vehicle.

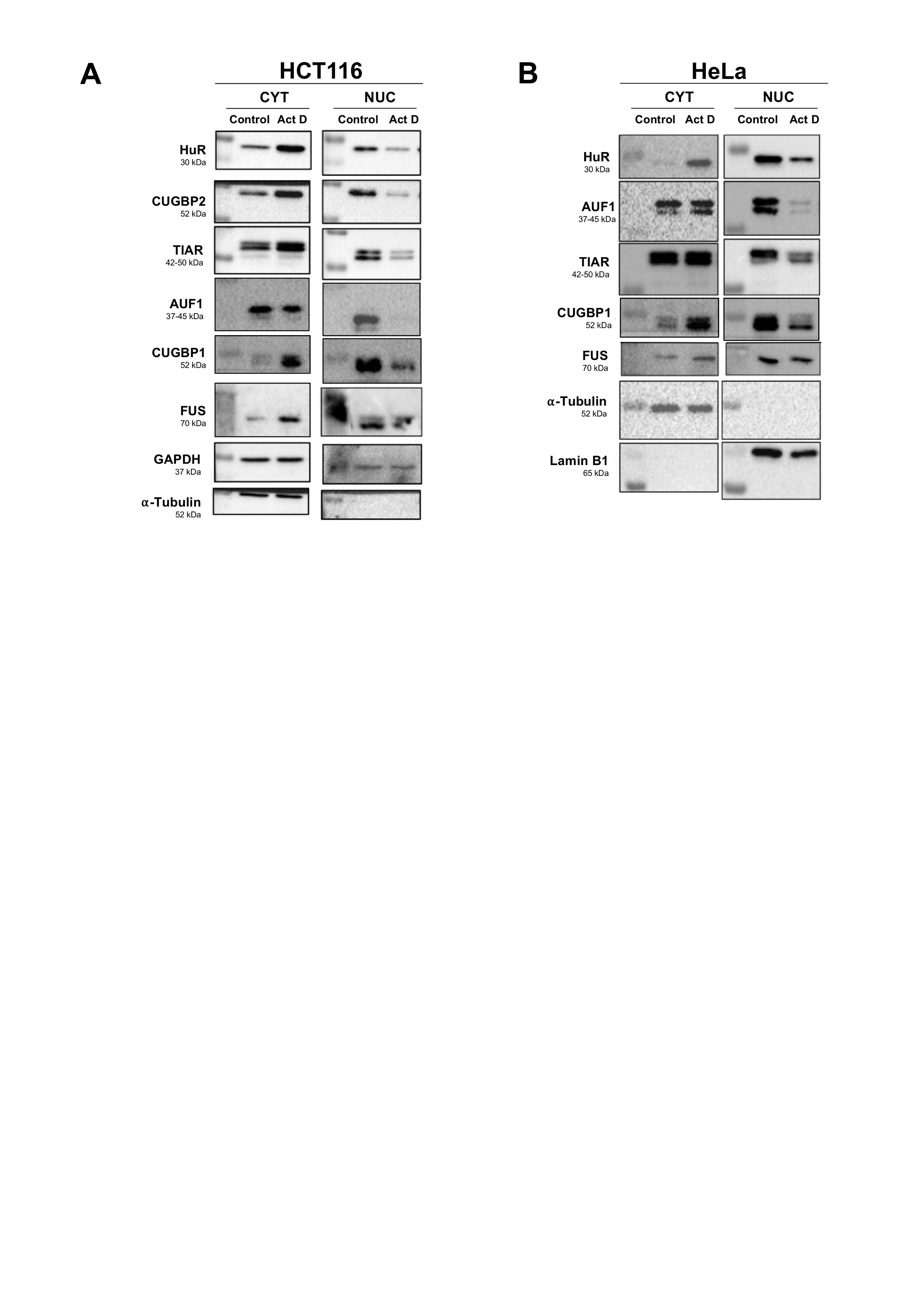

**Supplementary Figure 12. FUS and other RBPs translocated to the cytoplasm upon Act D treatment.** Effect of 3 h Act D treatment on the subcellular localization of multiple RBPs were examined in HCT116 (A) and HeLa (B) cells in cytoplasmic (CYT) and nuclear (NUC) fractions.

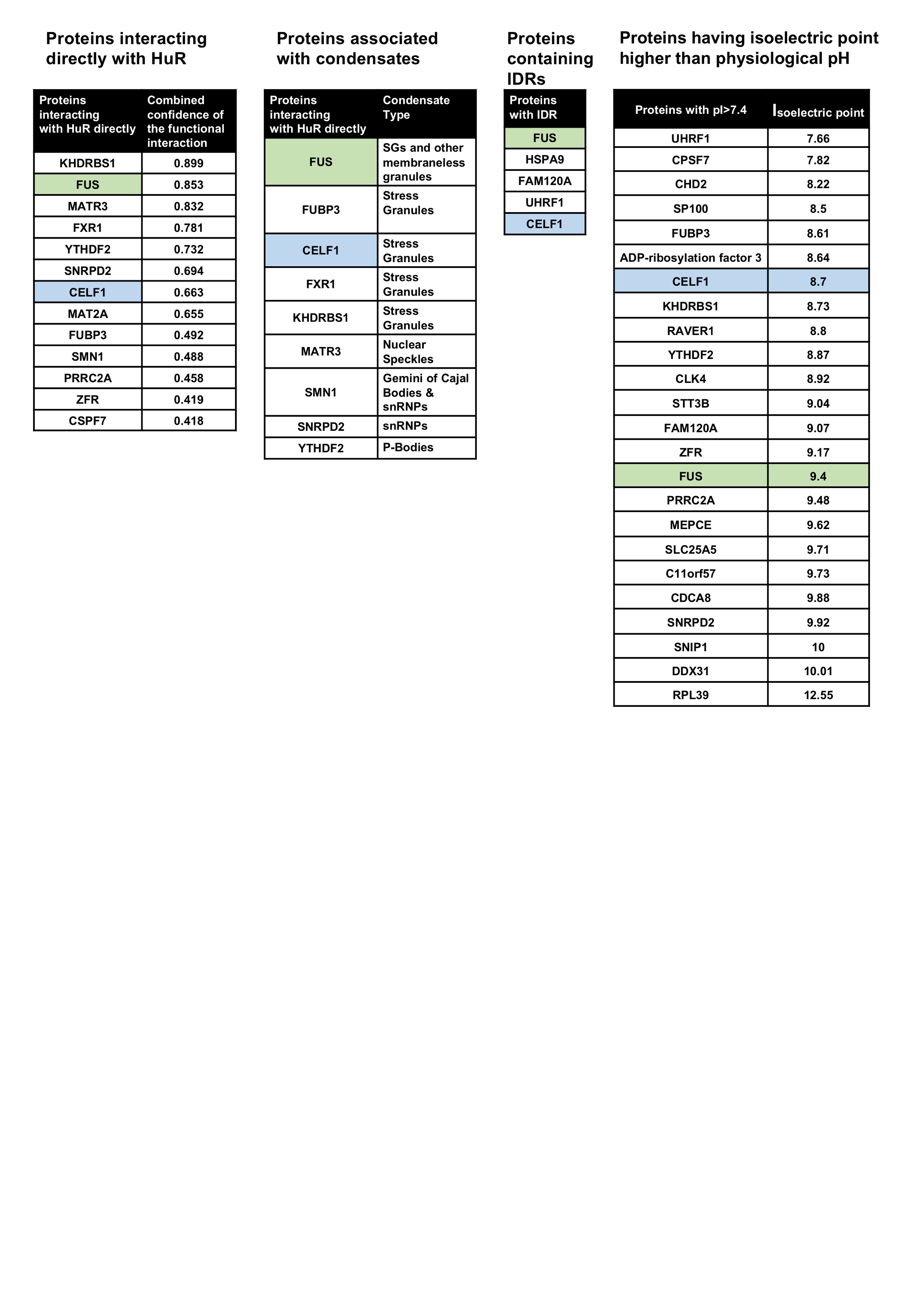

**Supplementary Figure 13. Proteins in close proximity to HuR were categorized to identify the potential interaction partner of HuR.** Candidate proteins were analyzed based on four criteria: Reported direct interaction with HuR, Association with condensate-related structures, presence of intrinsically disordered region (IDR), and high isoelectric point (pI). STRING database and literature search were used for protein categorization

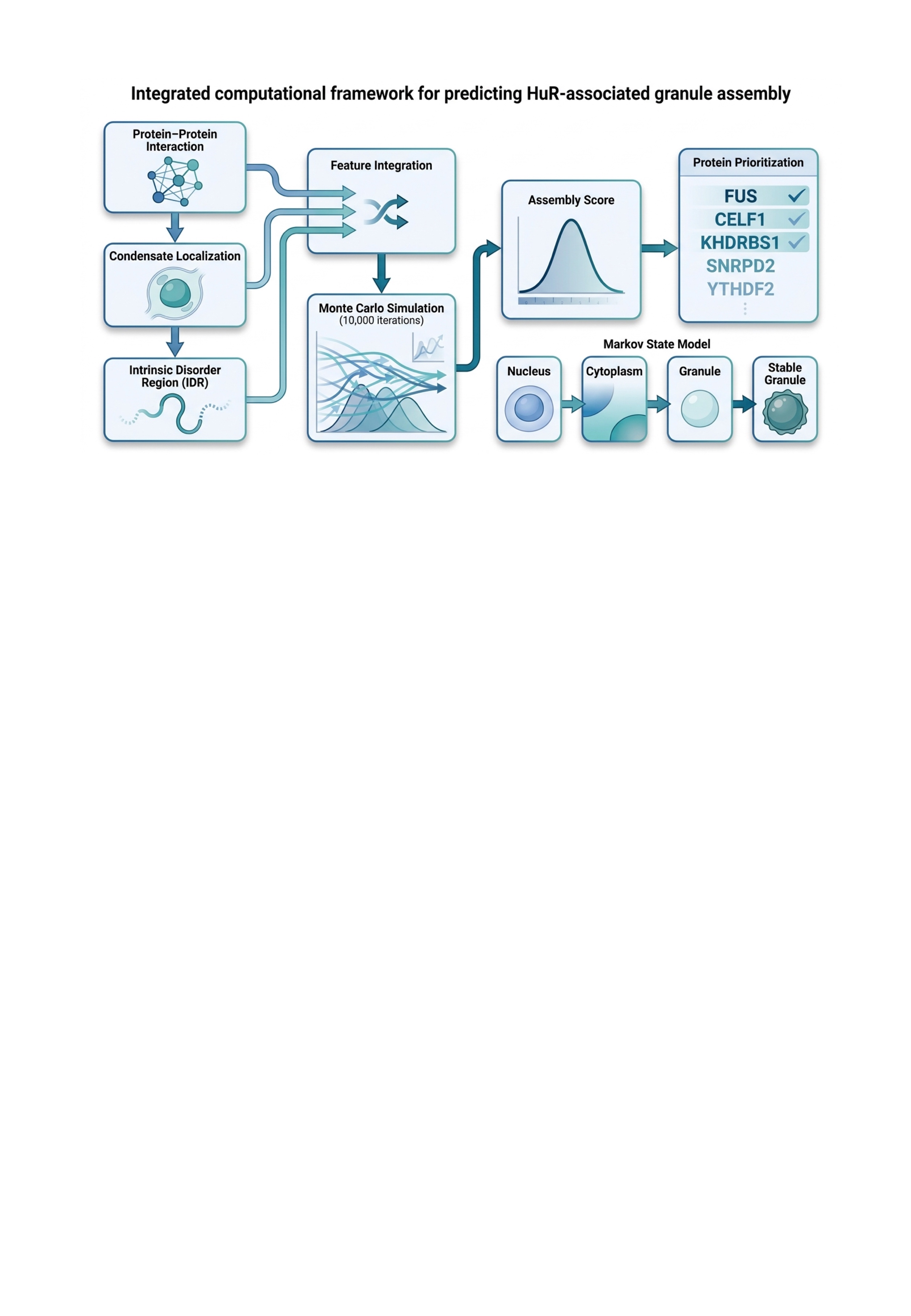

**Supplementary Figure 14. Integrated computational framework for predicting HuR-associated granule assembly.** Multiple experimental protein features (protein-protein interaction, condensate localization, and intrinsic disorder region) are integrated using a probabilistic model and Monte Carlo simulation to generate assembly scores, which guide Markov state modelling of granule formation and prioritize proteins most likely to be associated with HuR-based condensates.

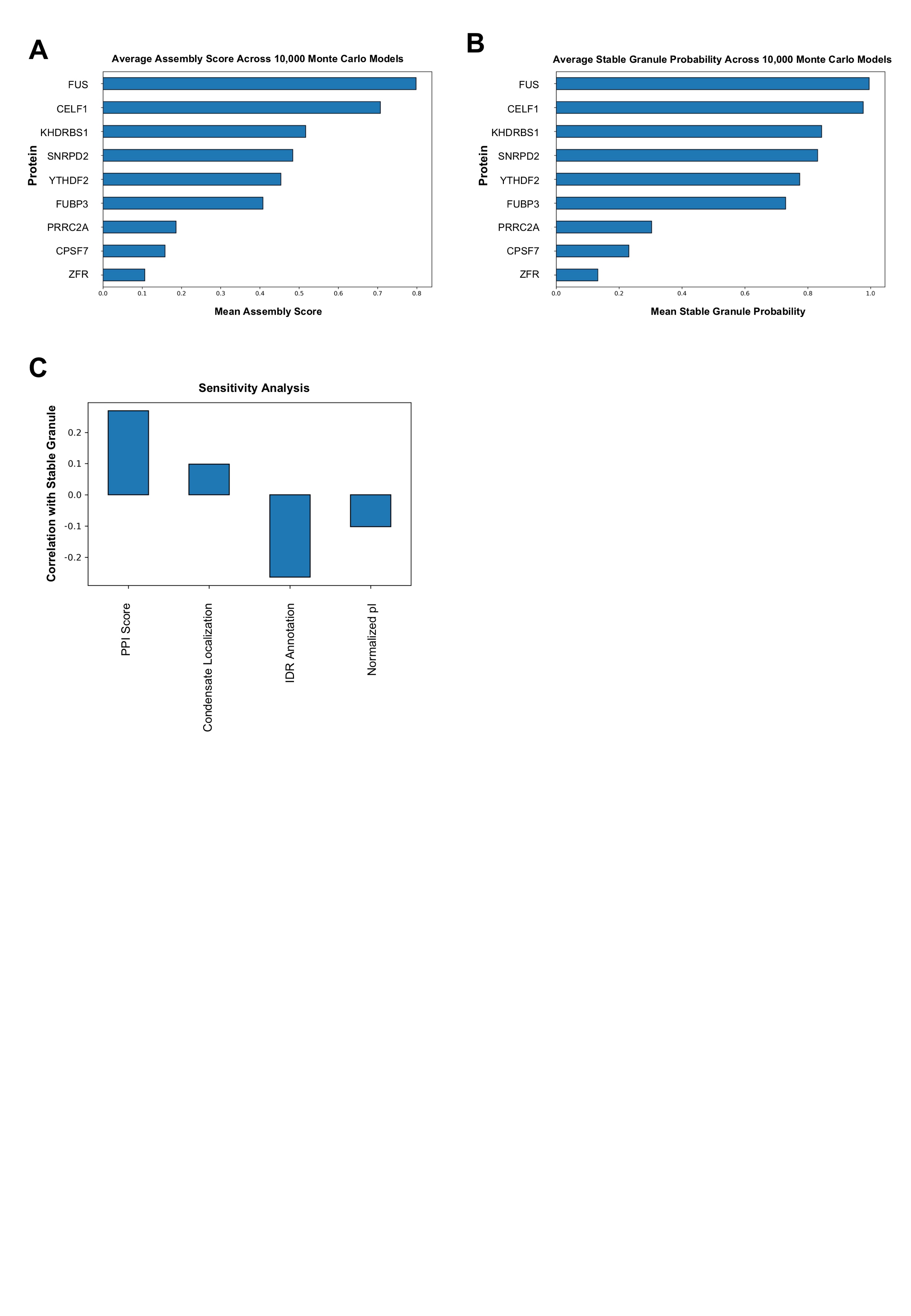

**Supplementary Figure 15. Probabilistic Modelling revealed that FUS was the protein with the highest propensity to form stable granules with HuR**. (A) Proteins were ranked according to their mean Assembly Score. Higher scores indicated greater predicted propensity for HuR-associated granule assembly. (B) Stable Granule probabilities were estimated using the discrete-time Markov model: Proteins with higher Assembly Scores consistently exhibited higher probabilities of reaching the Stable Granule state. (C) Correlations between sampled feature weights and predicted Stable Granule probabilities across Monte Carlo simulations were shown. Positive values indicated features that increased the likelihood of stable granule formation, whereas negative values indicated inverse associations within the current probabilistic model.

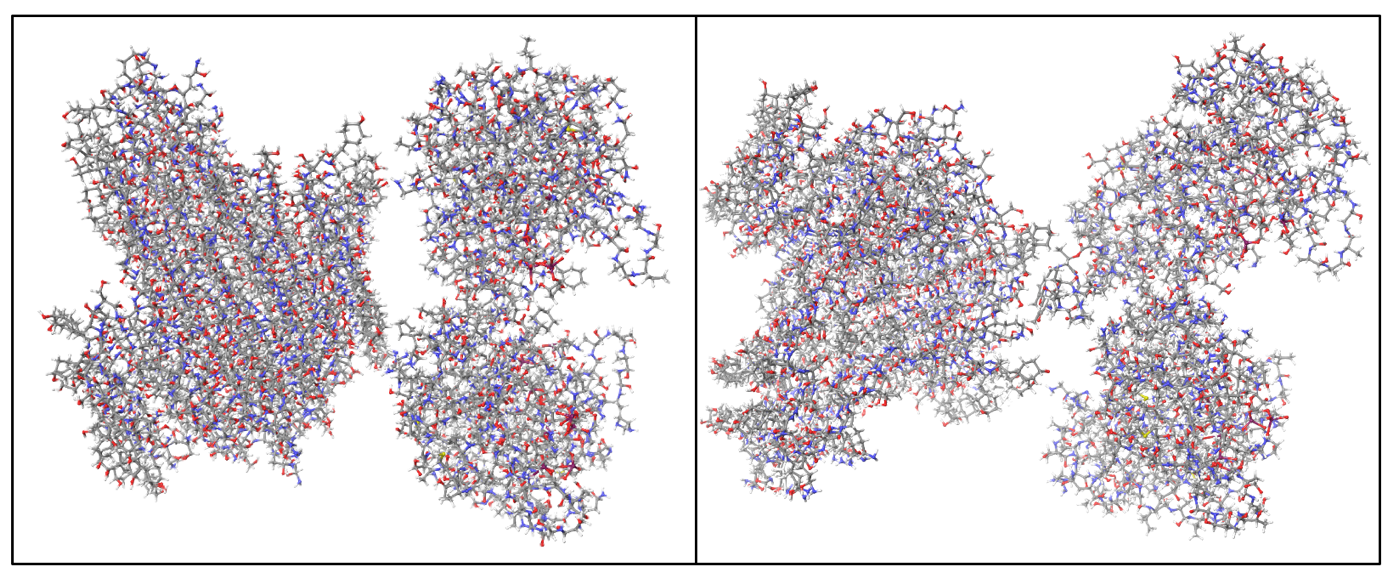

**Supplementary Figure 16. Addition of Act D molecules enhanced the interaction between HuR and FUS *in silico*.** Docking modelling suggested that the interaction between HuR and FUS can be stabilized by the presence of Act D molecules. The resulting docking model had an energy profile of -97.110 kcal/mol.

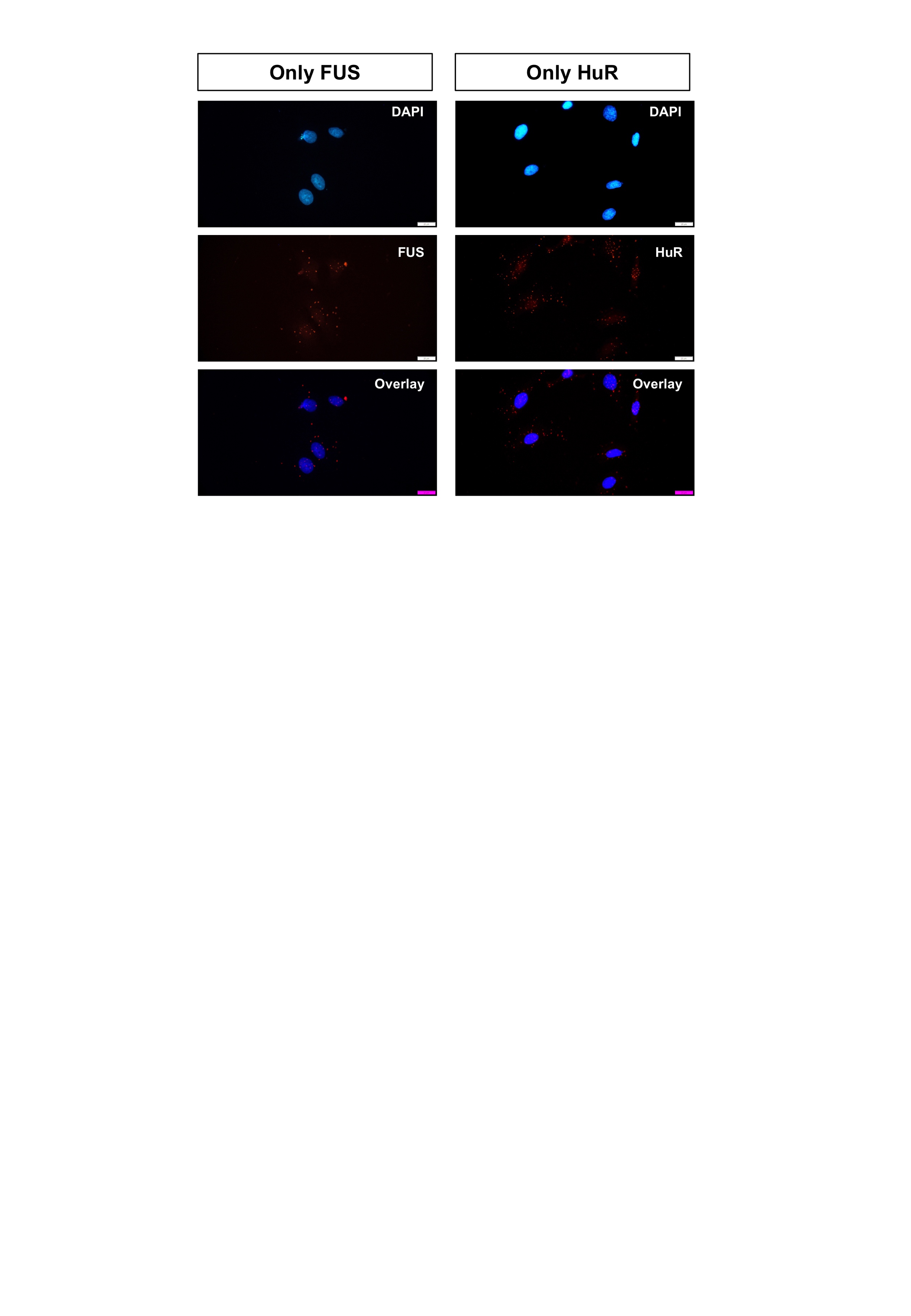

**Supplementary Figure 17. Control experiments revealed specificity of the enhanced interaction between HuR and FUS with Act D treatment**. In the proximity ligation assay (PLA), control groups included fixed cells that were incubated with FUS antibody alone or with HuR antibody alone to determine non-specific signals. Incubation with single antibody showed minimal number of red puncta. Images were acquired using Olympus BX43 microscope.

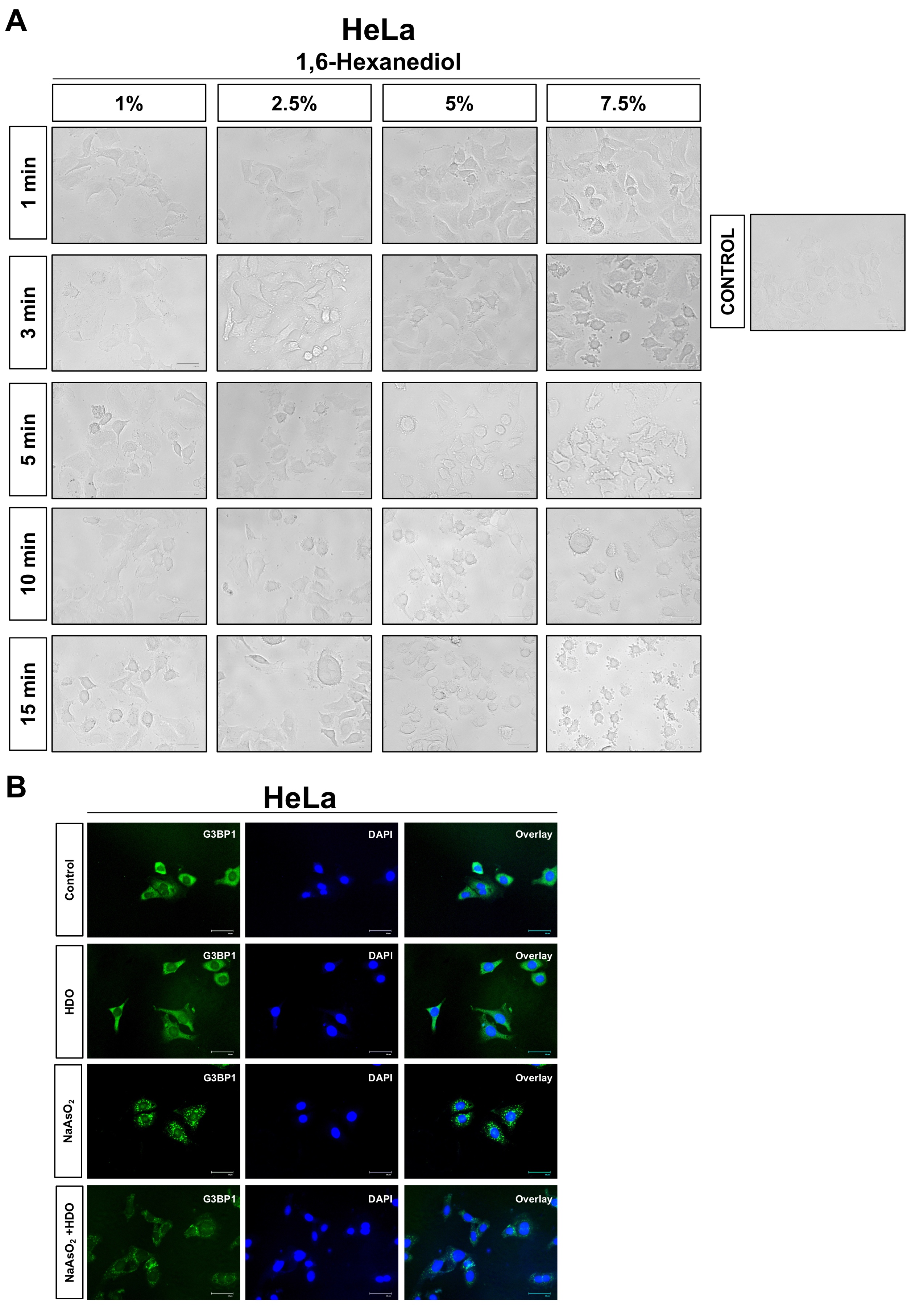

**Supplementary Figure 18. Optimal 1,6 Hexanediol treatment conditions did not disrupt Na-Arsenite-induced stress granules.** (A) Four different concentrations and five treatment durations were tested to observe the effects of 1,6-Hexanediol on cell morphology and viability. (B) Treatment with 2.5% 1,6-Hexanediol (HDO) for 3 min was selected as the optimal condition due to minimal cellular toxicity, however this concentration and duration did not reverse the effect of Na-Arsenite (NaAsO_2_) in the formation of stress granules (SGs). SGs were visualized by incubating the fixed cells with G3BP1 antibody followed by Alexafluor 488-conjugated secondary antibody. SGs were imaged using an EVOS™ Floid™ Cellular Imaging System.

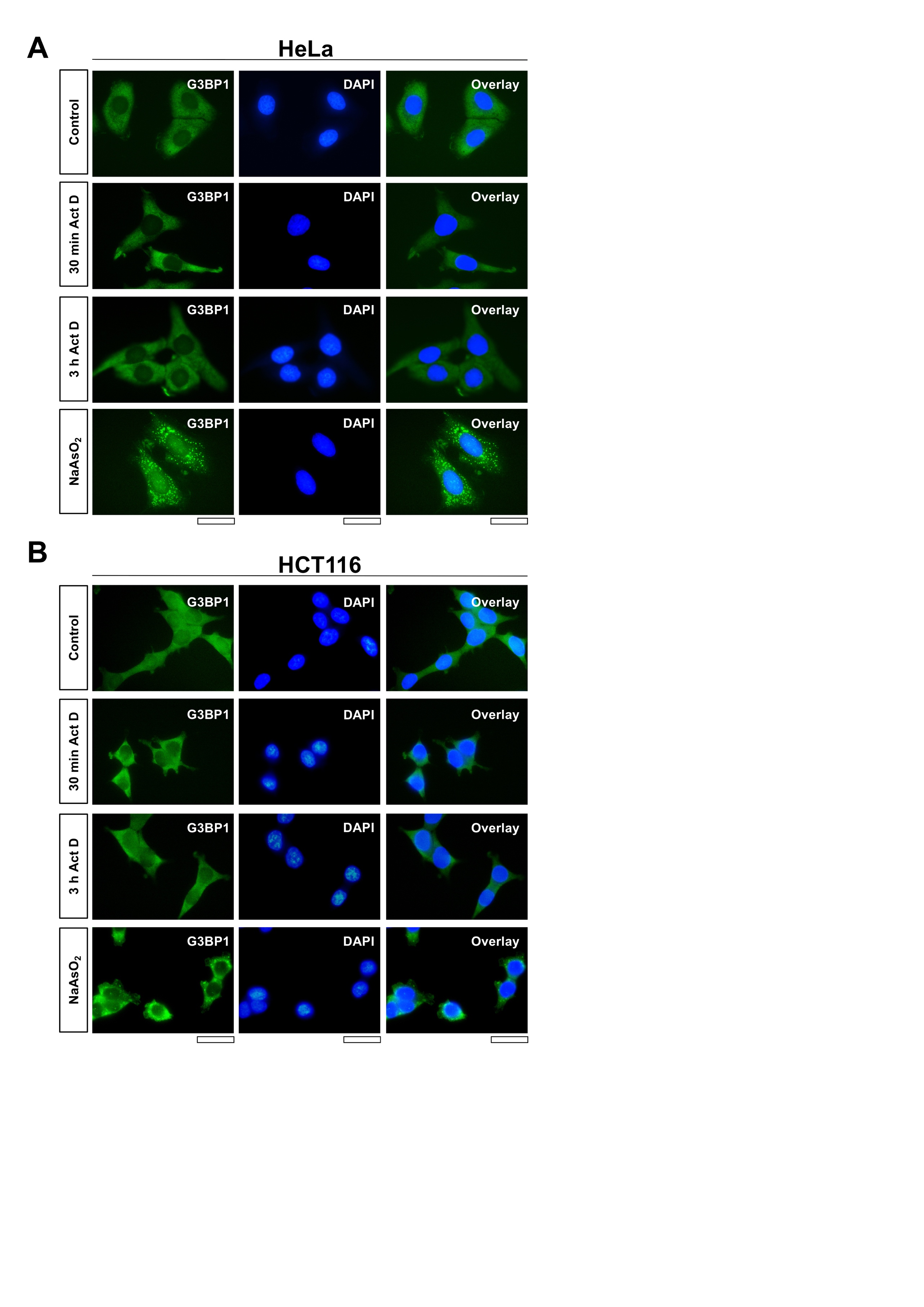

**Supplementary Figure 19. Act D treatment did not induce the formation of stress granules.** HeLa (A) and HCT116 (B) cells were treated with 10 μg/mL Act D for 30 min or 3 h, or with 200 μM Na-Arsenite (NaAsO_2_) for 1 h. The cells were subsequently fixed, permeabilized and incubated with G3BP1 antibody followed by AlexaFlour 488-conjugated secondary antibody. Finally, the nuclei were stained with DAPI containing mounting meidum. Stress Granules were detected only in Na-Ars treated cells (bottom panels). Images were acquired using an EVOS™ Floid™ Cellular Imaging System.

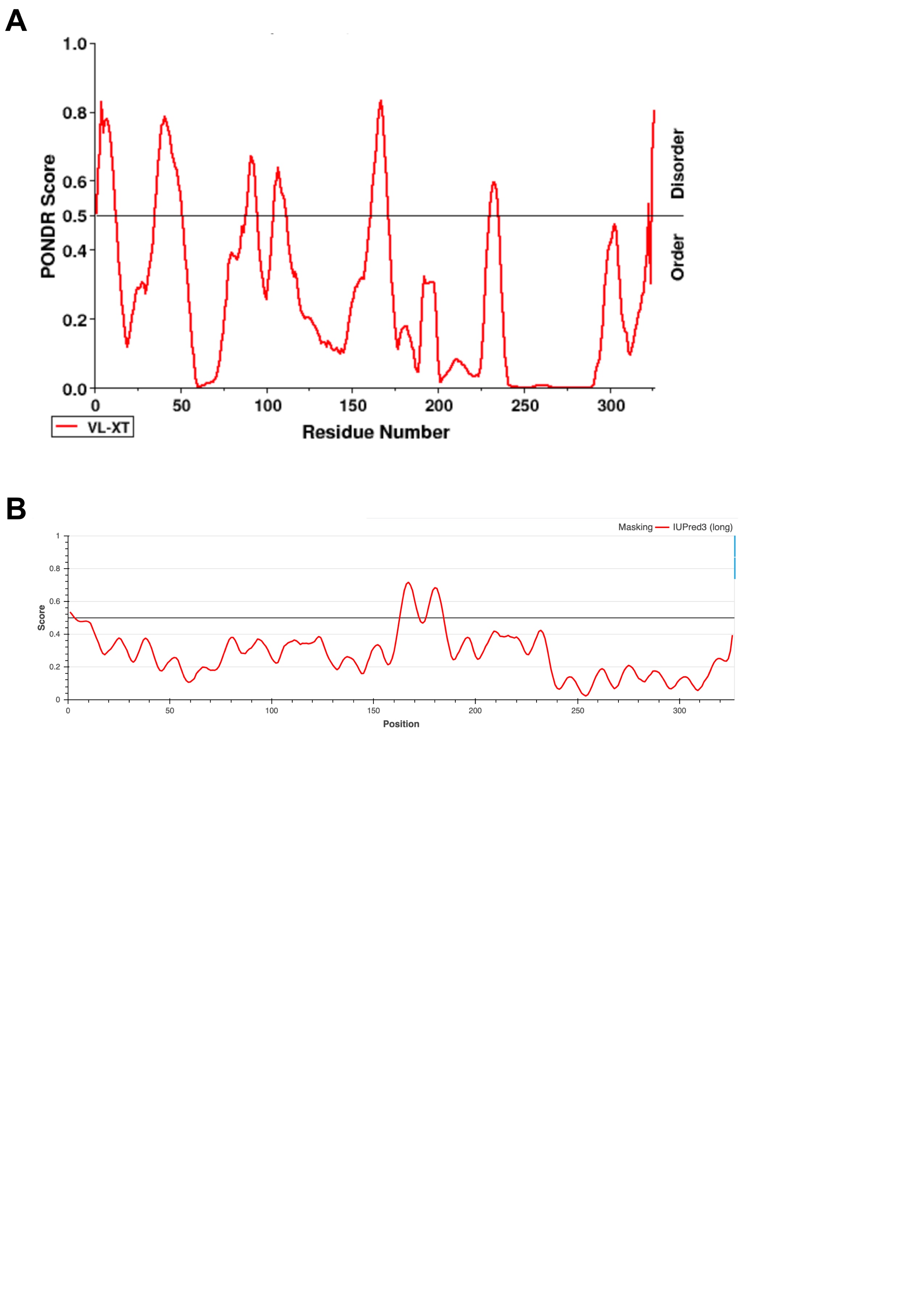

**Supplementary Figure 20. Prediction analysis of intrinsically disordered region of HuR showed that the protein has both structured and disordered regions**. Intrinsic disorder analysis of HuR using PONDR (A) indicated multiple intrinsically disordered regions distributed throughout the HuR protein sequence, with prominent disorder peaks at the N-terminus, central region (~150–175), and C-terminus. In contrast, IUPred3 (B), which applies a more stringent prediction algorithm, identified a disordered region in the central part of the protein while showing most of the other regions as structured.
